# Mutations in Parkinsonism-linked endocytic proteins synaptojanin1 and auxilin have synergistic effects on dopaminergic axonal pathology

**DOI:** 10.1101/2022.02.28.482419

**Authors:** Xin Yi Ng, Yumei Wu, Youneng Lin, Sidra Mohamed Yaqoob, Lois E. Greene, Pietro De Camilli, Mian Cao

**Affiliations:** Programme in Neuroscience and Behavioural Disorders, Duke-NUS Medical School, Singapore; Department of Physiology, National University of Singapore, Singapore; Departments of Neuroscience and Cell Biology, Howard Hughes Medical Institute, Program in Cellular Neuroscience, Neurodegeneration and Repair, Kavli Institute for Neuroscience, Yale University School of Medicine, New Haven, CT 06510, USA. Aligning Science Across Parkinson’s (ASAP) Collaborative Research Network, Chevy Chase, MD; Laboratory of Cell Biology, NHLBI, National Institutes of Health, MD, USA

**Author notes:** These authors contributed equally (Yumei Wu, Youneng Lin, Sidra Mohamed Yaqoob). Other authors’ emails: Xin Yi Ng Yumei Wu Youneng Lin Sidra Mohamed Yaqoob Lois E. Greene Pietro De Camilli.

**Keywords:** Synaptojanin1, clathrin, DAergic terminals, Parkinsonism, DNAJC6/PARK19, SYNJ1/PARK20, PI4P, PI(4,5)P_2_, inositol-phosphatase, GAK

## Abstract

Parkinson’s disease (PD) is a neurodegenerative disorder characterized by defective dopaminergic (DAergic) input to the striatum. Mutations in two genes encoding synaptically enriched clathrin-uncoating factors, synaptojanin 1 (SJ1) and auxilin, have been implicated in atypical Parkinsonism. SJ1 knock-in (SJ1-KI^RQ^) mice carrying a disease-linked mutation display neurological manifestations reminiscent of Parkinsonism. Here we report that auxilin knockout (Aux-KO) mice display dystrophic changes of a subset of nigrostriatal DAergic terminals similar to those of SJ1-KI^RQ^ mice. Furthermore, Aux-KO/SJ1-KI^RQ^ double mutant mice have shorter lifespan and more severe synaptic defects than single mutant mice. These include increase in dystrophic striatal nerve terminals positive for DAergic markers and for the PD risk protein SV2C, as well as adaptive changes in striatal interneurons. The synergistic effect of the two mutations demonstrates a special lability of DAergic neurons to defects in clathrin uncoating, with implications for PD pathogenesis in at least some forms of this condition.

## Introduction

Recessive loss-of-function (LOF) mutations in two genes *SYNJ1 (PARK20)* and *DNAJC6 (PARK19),* which encode the pre-synaptically enriched proteins synaptojanin 1 (SJ1) and auxilin, were reported in rare cases of familial recessive juvenile/early-onset atypical Parkinsonism ^1–4^. Interestingly, these two proteins function at different steps, albeit synergistically, in the shedding of the clathrin coat that follows clathrin-dependent vesicle budding to generate new synaptic vesicles (SVs) (Fig. 2A) ^5^. SJ1, a phosphatase that sequentially dephosphorylates PI(4,5)P_2_ via tandemly arranged 5-phosphatase and 4-phosphatase domains, mediates the dissociation of the endocytic clathrin adaptors (the inner layer of the coat), whose membrane interaction is PI(4,5)P_2_ dependent ^6^. Auxilin, a clathrin-binding co-chaperone, recruits the ATPase HSC70 to disassemble the clathrin lattice, i.e. outer layer of the coat ^6^. Moreover, since auxilin contains a binding domain for monophosphoinositides ^7^, its recruitment was proposed to be controlled, at least in part by the activity of SJ1.

Complete LOF of SJ1 results in early postnatal death ^8, 9^, while the R258Q mutation responsible for Parkinsonism selectively impairs the function of its 4-phosphatase domain (also called Sac domain) ^2^. In the case of auxilin, Parkinsonism mutations result in complete or very strong loss of function ^10^. Patients affected by Parkinsonism carrying mutations in these two genes share similar clinical manifestations: early- or juvenile-onset, typical motor deficits with atypical epilepsy and, in some patients, developmental delay and/or DA deficiency as detected by reduced DaTScan signal in the striatum ^11, 12^.

Several animal models have been generated and characterized for both Parkinsonism genes. Knock-in (KI) mice carrying the Sac domain mutation R258Q (R259Q in mice, referred henceforth as SJ1-KI^RQ^ mice), recapitulated patients’ neurological manifestations ^13^. Nerve terminals of these mice exhibited accumulation of clathrin coated vesicles (CCVs) and SVs endocytic defects. Importantly, selective synaptic abnormalities were observed in nigrostriatal DAergic nerve terminals of these mice, i.e. the axonal projections specifically degenerated in PD ^13^. Perturbations in autophagosome formation were also reported in neuronal cultures of these mice ^14^, in patient iPSC-derived human DA neuronal models carrying the same mutation and in nerve terminals of *Drosophila* carrying the corresponding mutation ^15^. Motor symptoms, endocytic defects at synapses and degeneration of DAergic nerve terminals were also reported in aged SJ1 Heterozygous (+/-) mice ^16^. Moreover, a strong synthetic interaction was observed in mice between the SJ1 R258Q mutation and the loss of Sac2, a PI4P phosphatase shown by GWAS to be a candidate PD risk gene ^17^.

LOF animal models of auxilin also have features of PD pathogenesis. Reduced auxilin level in *Drosophila* leads to PD-like motor symptoms and accelerates α-synuclein mediated DA neuron loss ^18^. Auxilin mutant flies are also more sensitive to the environmental toxin paraquat^18^. Auxilin KO mice (Aux-KO mice) are born with a lower than normal mendelian ratio and a subset of them die perinatally ^19^. The brain levels of the auxilin paralogue, cyclin G-associated kinase (GAK), are upregulated in the surviving mice, and these mice were reported to have a normal lifespan, although a systematic analysis of their neurological performance was not carried out. In neuronal cultures derived from these mice, impaired SVs endocytosis and clathrin uncoating defects were observed ^19^. Mice carrying the auxilin Parkinsonism-linked mutation R927G showed motor impairments in old mice, and both SVs recycling and Golgi trafficking defects ^20^. Finally, in human midbrain-like organoids, mutations of auxilin were shown to cause key PD pathologic features and also developmental defects due to impaired WNT-LMX1A signaling ^21^.

The implication in Parkinsonism of two genes that have cooperative function in clathrin uncoating during SVs recycling strongly suggest that their mutations result in Parkinsonism via the same, or very similar, pathogenetic mechanism. Elucidating such a mechanism may offer the possibility of developing intervention strategies to prevent the onset of Parkinsonism manifestations in individuals homozygous for these mutations. It is also possible that mutations of other PD genes may converge on such mechanisms, so that studies on SJ1 and auxilin may have more general implications.

Here we set out to investigate whether loss of auxilin function in mice phenocopies the dystrophic changes that we have observed in the striata of SJ1-KI^RQ^ mice and to determine whether the combined loss of auxilin and the Parkinsonism mutation of SJ1 results in synergistic effects on such changes. Our results demonstrate a similar effect of the two genetic perturbations on nigrostriatal DAergic nerve terminals. We also show that Aux-KO/SJ1-KI^RQ^ double mutant mice have a much shorter lifespan than single mutants and more severe dystrophic changes in the striatum. Our results strengthen evidence for a vulnerability of nigrostriatal neurons to clathrin uncoating perturbations at synapses, and suggest that such dysfunction may play a role in at least some form of Parkinsonism.

## Results

### Neurological defects in Aux-KO mice that survive early postnatal death

As previously reported, a subset of Aux-KO mice die shortly after birth ^19^. The survivors can have a lifespan similar to wild-type (WT) controls but a subset of them develop early-onset tonic-clonic epileptic seizures starting at 3-4 postnatal week (Supplementary Mov. 1), a finding also reported in the clinical cases carrying auxilin mutations. Hindlimb clasping phenotype, a sign of neurodegeneration, was observed in 30% of the mice at 1-month-old (Fig. 1A). Moreover, at 2-4 months they showed a mild fine motor deficit in the balanced beam test with significantly more missteps as they walk along the beam (Fig. 1B, C and Supplementary Mov. 2A, 2B), while their performance on rotarod (gross motor) was normal (Fig. 1D). These results show that Aux-KO mice have at least some of the features observed in patients carrying auxilin Parkinsonism mutations.

**Figure 1.**
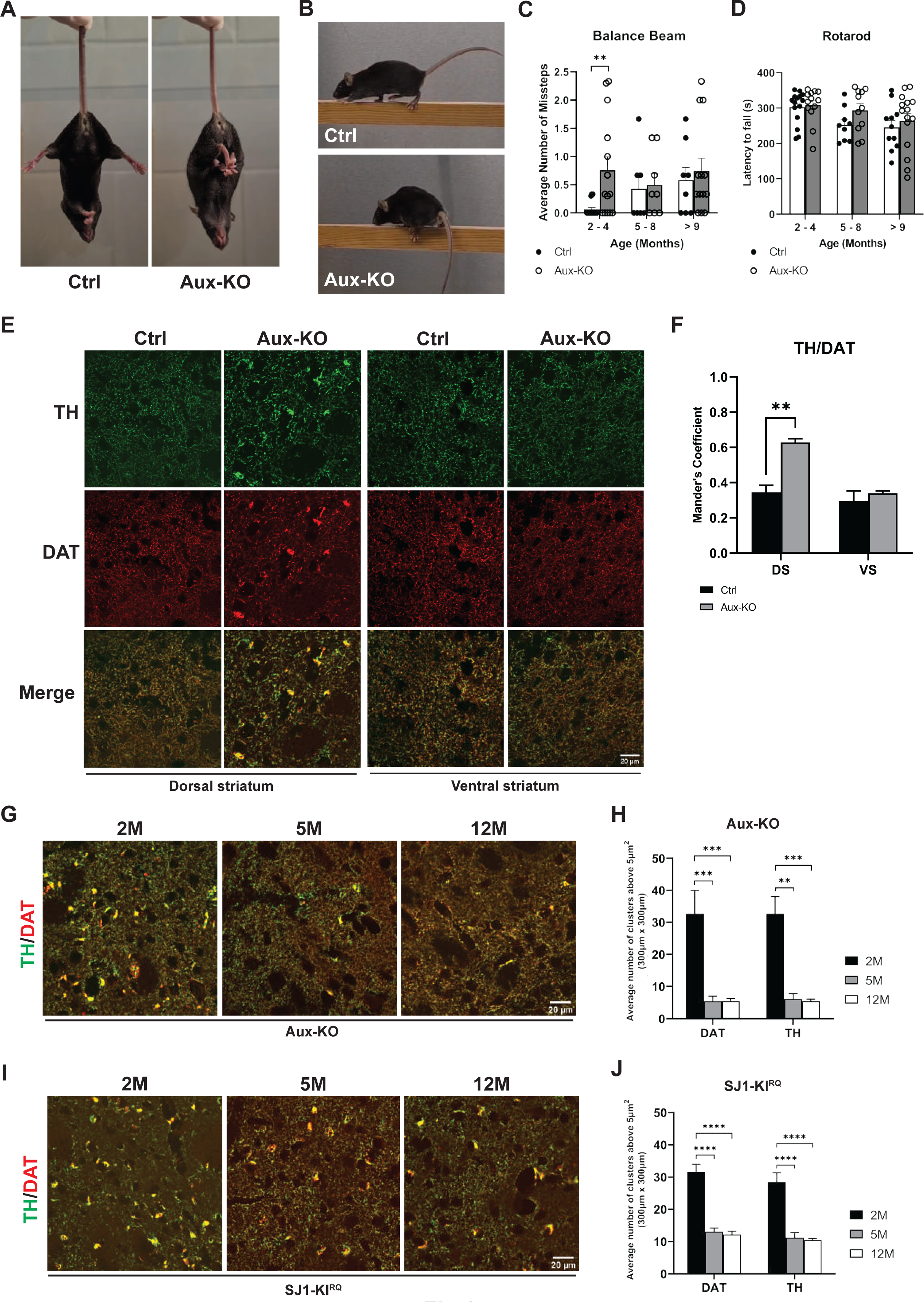
Aux-KO mice exhibit parkinsonism-like phenotypes with dystrophic DAergic axon terminals in the dorsal striatum. (A) Hindlimb clasping phenotype of 4-month-old Aux-KO mouse. (B) 2-month-old Aux-KO mice have missteps while walking on the balance beam. (C) Quantification for the number of missteps during the balance beam test for three age groups of Aux-KO mice. Aux-KO mice only exhibit mild fine motor deficits at 2-4 months compared to the Ctrl group. Data are represented as mean ± SEM (** *p*<0.01, by Student’s unpaired *t* test). N numbers for the Ctrl (2-4M: 15, 5-8M: 7 and >9M: 8) and Aux-KO mice (2-4M: 14, 5-8M: 8 and >9M: 13). (D) Performance of three age groups of Aux-KO mice on the accelerated rotarod. Data are represented as mean ± SEM. N numbers used for each age group of the Ctrl and Aux-KO mice (2-4M: 15 and 13; 5-8M: 9 and 11; >9M: 11 and 15). (E) Double staining for TH and DAT of 1-month-old Ctrl and Aux-KO mice shows selective clustering of these two DAergic markers only in the dorsal striatum. (F) Mander’s colocalization coefficient showed that TH and DAT clusters which are only found in the dorsal striatum (DS) colocalizes very well with each other. Data are represented as mean ± SEM (** *p*<0.01, by unpaired t test). N = 3 mice for each genotype. (G and I) Age-dependent decrease of the number of TH/DAT-positive clusters in the dorsal striatum of Aux-KO and SJ1-KI^RQ^ mice is observed respectively at both 5-month and 12-month. (H and J) Quantification for TH and DAT clusters shown in (G) and (I). The number of clusters is counted in five randomly selected, 300 x 300-µm regions of interest (ROIs). Data are represented as mean ± SEM (** *p*<0.01, and *** *p*<0.001, and **** *p*<0.0001 by two-way ANOVA with post-hoc Tukey’s test). n = 3 mice for each genotype and each age group.

### Defects of DAergic nerve terminals in the dorsal striatum of Aux-KO mice

The previously reported analysis of cultured neurons of Aux-KO mice demonstrated perturbations consistent with a defect in clathrin uncoating. To examine the impact of auxilin LOF on the DAergic nigrostriatal pathway, we performed immunohistochemistry on brain frozen sections using antibodies directed against tyrosine hydroxylase (TH) and the plasma membrane dopamine transporter (DAT), both markers of DA neurons. Similar to SJ1-KI^RQ^ mice, scattered abnormal TH- and DAT-positive clusters, were observed in the striata of these mice, but not in WT and heterozygous littermates, in addition to the normal diffuse punctate immunoreactivity representing axon terminal varicosities of DAergic neurons (Fig. 1E, F). Typically, these clusters were localized next to the soma of striatal neurons. Interestingly, as in the case of SJ1-KI^RQ^ brains, these clusters were specifically present in the dorsal but not ventral striatum (Fig. 1E). Thus, nerve terminals of the nigrostriatal DA system are selectively and similarly affected by either complete LOF of auxilin or the R258Q mutation in SJ1, two genetic perturbations that result in Parkinsonsim in human patients. Tiny DAT-positive clusters first appeared at about 3-week and large clusters positive for both DAT and TH peaked at 1-2 months. However, in both adult (5-month) and aged mice (12-month), the number of these clusters was significantly reduced (Fig. 1G, H), similar to what we observed in SJ1-KI^RQ^ mice (Fig. 1I, J). Possibly, structural changes of a subset of DAergic nerve terminals only occurs at a young age and these terminals eventually degenerate. However, the occurrence of a repair mechanism cannot be excluded.

### More severe neurological phenotypes in Aux-KO/SJ1-KI^RQ^ double mutant mice than in single mutant mice

The similar clinical phenotypes of Parkinsonism mutations in SJ1 (RQ mutation) and auxilin (complete LOF) as well as the similar cellular and histological phenotypes observed in SJ1-KI^RQ^ and Aux-KO mice prompted us to examine a synergistic effect of the two genetic disruptions by generating Aux-KO/SJ1-KI^RQ^ double mutant mice. As shown in Fig. 2B, Aux-KO/SJ1-KI^RQ^ mice have much shorter lifespan compared to either single mutant mice, revealing a strong synthetic genetic interaction. About 75% of Aux-KO/SJ1-KI^RQ^ mice died within 1 month after birth. Moreover, they were usually smaller in size (Supplementary Fig. 1A, B) and showed hindlimb clasping and seizures. The level of Neuropeptide Y (NPY) was dramatically increased in dentate gyrus (DG) mossy fibers in Aux-KO/SJ1-KI^RQ.^ hippocampus, compared to single mutants and controls, most likely a sign of frequent epileptic seizures ^22, 23^ (Supplementary Fig. 1C, D). To date, only one Aux-KO/SJ1-KI^RQ^ mouse out of more than 50 survived to adulthood and died suddenly at 7-month-old. As assessed by the balanced beam test and by immunohistological analysis, this mouse had more severe motor coordination defect and DAergic terminal dystrophy in the striatum compared to single mutant mice (Supplementary Mov. 3A-D and Supplementary Fig. 1E) Hematoxylin and eosin (H&E) staining of Aux-KO/SJ1-KI^RQ^ mouse brains at 1-month of age (the age at which nearly 25% of the double mutant mice survived) revealed a normal brain architecture (Supplementary Fig. 2). Likewise, no obvious gliosis was observed in different brain regions upon immunostaining with ionized calcium-binding adapter molecule (Iba1), a microglia marker and glial fibrillary acidic protein (GFAP), an astrocyte marker (Supplementary Fig. 3A, B). Moreover, western blot analysis of total homogenates of brains at the same age showed that the levels of major synaptic proteins tested, including SV proteins, endocytic proteins and proteins of DA metabolism were unchanged at this age, with the exception of a modest upregulation in amphiphysin 2 and endophilin1 of SJ1-KI^RQ^ mice (Supplementary Fig. 4A, B).

**Figure 2.**
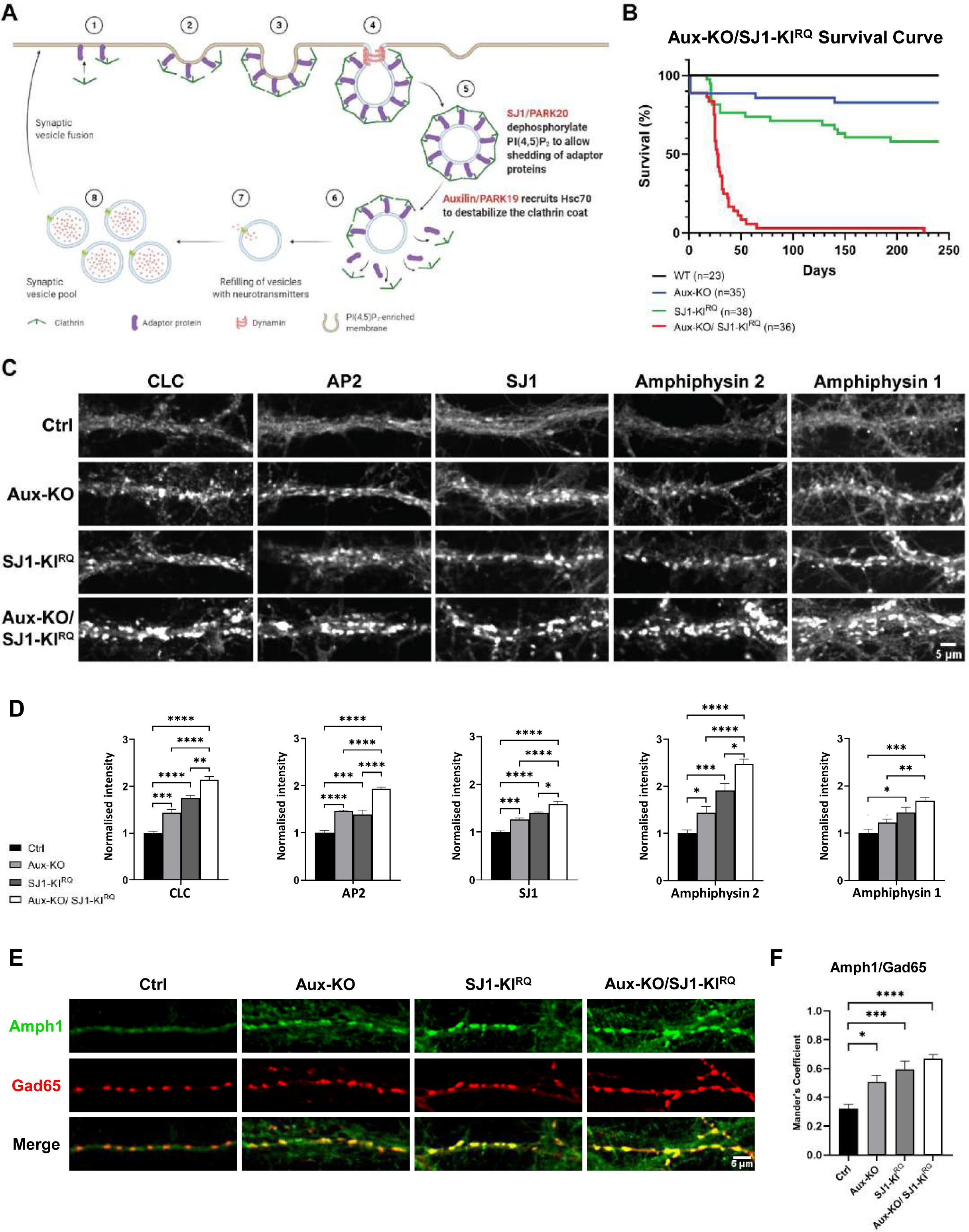
LOF of both Auxilin and SJ1 exacerbate neurological defects in Aux-KO/SJ1-KI^RQ^ mice. (A) Schematic diagram displaying the role of Auxilin (PARK19) and SJ1 (PARK20) in clathrin-mediated SVs recycling. (B) Survival curves of wild-type (WT), Aux-KO, SJ1-KI^RQ^, and Aux-KO/SJ1-KI^RQ^ mice. (C) Representative images of immunoreactivity for CLC, AP2, SJ1, Amphiphysin 2 and Amphiphysin 1 in DIV19 cultured primary cortical neurons from Ctrl, Aux-KO, SJ1-KI^RQ^ and Aux-KO/SJ1-KI^RQ^ newborn pups. (D) Quantification of synaptic clusters intensity shown in (C). Data are represented as mean ± SEM (* *p*<0.05, ** *p*<0.01, *** *p*<0.001, **** *p*<0.0001 by one-way ANOVA with post-hoc Tukey’s test). The number of neurons quantified for each endocytic marker is as follow: Ctrl: n=43-49, Aux-KO: n=32-52, SJ1-KI^RQ^: n=35-37, Aux-KO/SJ1-KI^RQ^: n=39-44 cultured from 4-6 mice. (E) Immunostaining of amphiphysin 1 with Gad65 (inhibitory presynaptic marker) in DIV19 cortical neurons revealed that these synaptic clustering of endocytic proteins occur predominantly in GABAergic neurons. (F) Mander’s coefficient shows that both single mutants (Aux-KO and SJ1-KI^RQ^) and double mutant Aux-KO/SJ1-KI^RQ^ have better colocalization of Amph1 and Gad65 clusters compared to control. Data are represented as mean ± SEM (* *p*<0.05, *** *p*<0.001; **** *p*<0.0001 by one-way ANOVA with post-hoc Tukey’s test). N=8-10 neurons.

### Synergistic disrupting effects of auxilin and SJ1 mutations on clathrin coat dynamics in nerve terminals

Synapses of either Aux-KO or SJ1-KI^RQ^ mice display abnormal presynaptic clustering of endocytic factors, as revealed by immunofluorescence (Fig. 2C, D), and an accumulation of CCVs as well as empty clathrin cages in the case of Aux-KO neurons, as revealed by electron microscopy (EM) (Fig. 3). These defects, which result in defective endocytic recycling of SVs, are not restricted to DAergic nerve terminals, in agreement with the ubiquitous expression of SJ1 and auxilin in neurons. Moreover, they are generally more prominent at inhibitory nerve terminals as labeled by GAD65, which have higher tonic levels of activity (Fig. 2E, F). Both these defects were more severe at synapses of double mutant neurons. At these synapses, the fluorescence intensity of clusters of endocytic proteins, such as clathrin, adaptor protein 2 (AP2), SJ1, amphiphysin 1 and amphiphysin 2, which was already higher in Aux-KO and SJ1-KI^RQ^ single mutant neurons than in controls, was even higher in Aux-KO/SJ1-KI^RQ^ neurons (Fig. 2C, D).

**Figure 3.**
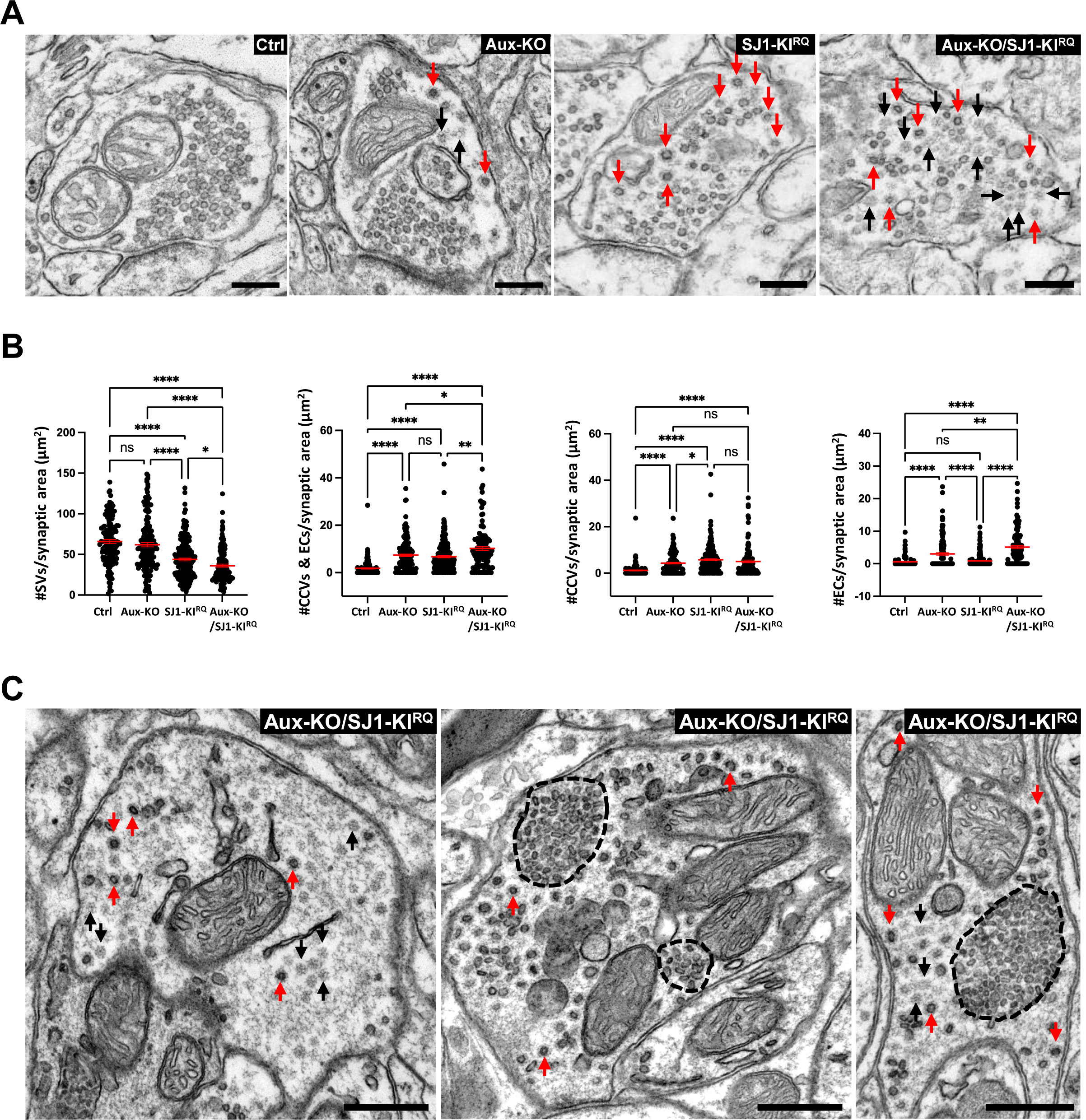
Accumulation of CCVs and empty clathrin cages in nerve terminals of Aux-KO, SJ1-KI^RQ^, and Aux-KO/SJ1-KI^RQ^ mice. (A) Representative EM micrographs of nerve terminals from the dorsal striatum of single and double mutant mice. Red and black arrows point to CCVs and empty clathrin coated cages (EC) respectively. Scale bar: 250 nm. (B) Morphometry analysis of the number of SVs, sum of number of CCVs and ECs, CCVs, as well as ECs per pre-synaptic area in the dorsal striatum (Ctrl n=124, Aux-KO n=156, SJ1-KI^RQ^ n=198, and Aux-KO/SJ1-KI^RQ^ n=120). Each dot represents one nerve terminal. Data are represented as mean ± SEM. * *p*<0.05, ** *p*<0.01, **** *p*<0.0001 (One-way ANOVA with Games-Howell’s multiple comparison test). (C) Examples of Purkinje nerve terminals in the deep cerebellar nuclei of Aux-KO/SJ1-KI^RQ^ mouse showing the striking accumulation of assembled clathrin (both ECs and CCVs). The tight packaging of SVs (middle and right images) was previously reported in nerve terminals of SJ1-KI^RQ^ mice (^13^ Fig. S3D) and may reflect dystrophic changes, Scale bar: 500 nm.

A synergistic effect of the two mutations could also be observed by EM, as shown by representative EM micrographs in Fig. 3A, and by a morphometric analysis of CCVs and empty clathrin cages in nerve terminals of the striatum (Fig. 3B). As expected, the SJ1-KI^RQ^ mutation resulted primarily in an increase in CCVs and the Aux-KO mutation also resulted in an increase of empty clathrin cages, but the combination of the two mutations led to an overall increase in assembled clathrin (both cages and coated vesicles). At some inhibitory synapses in other regions of the brain such increase was huge, as illustrated by the three examples of inhibitory nerve terminals in the deep cerebellar nuclei shown in Fig. 3C that show massive accumulations of empty clathrin cages and CCVs.

### Enhanced dystrophic changes in the striatum of Aux-KO/SJ1-KI^RQ^ double mutant mice

We next focused on the DA system in both midbrain and striatum. The morphology of neurons positive for TH and for aromatic L-amino acid decarboxylase (AADC, the enzyme that acts downstream of TH to convert levodopa into DA and is thus another marker of DAergic neurons) in both substantia nigra (SN) and ventral tegmental area (VTA) appeared normal in 1-month-old Aux-KO/SJ1-KI^RQ^ mice when compared to WT controls (Supplementary Fig. 5A). Stereological counting showed no significant loss of TH-positive neurons in either SN or VTA in Aux-KO/SJ1-KI^RQ^ midbrain at this age (Supplementary Fig. 5B). However, in the striatum of Aux-KO/SJ1-KI^RQ^ mice, the DAergic nerve terminal dystrophic phenotype was more severe than in the single mutant mice. The number of TH/DAT-positive clusters was significantly increased compared to single mutants (Fig. 4A, B, E). More importantly, similar TH/DAT-positive clusters were also observed in the ventral striatum (including both the nucleus accumbens and olfactory tubercle) of Aux-KO/SJ1-KI^RQ^ mice, but not of the single mutant mice (Fig. 4C, D, F and Supplementary Fig. 5C, 6). Since the ventral striatum receives DAergic axonal input from the VTA of the midbrain, this result indicates that the combined perturbation of auxilin and SJ1 affects the function of midbrain DAergic neurons more globally. It suggests a vulnerability of both SN and VTA DAergic neurons to the combined loss of SJ1 and auxilin, but a lower tolerance of SN neurons to the loss of either one of these two proteins.

**Figure 4.**
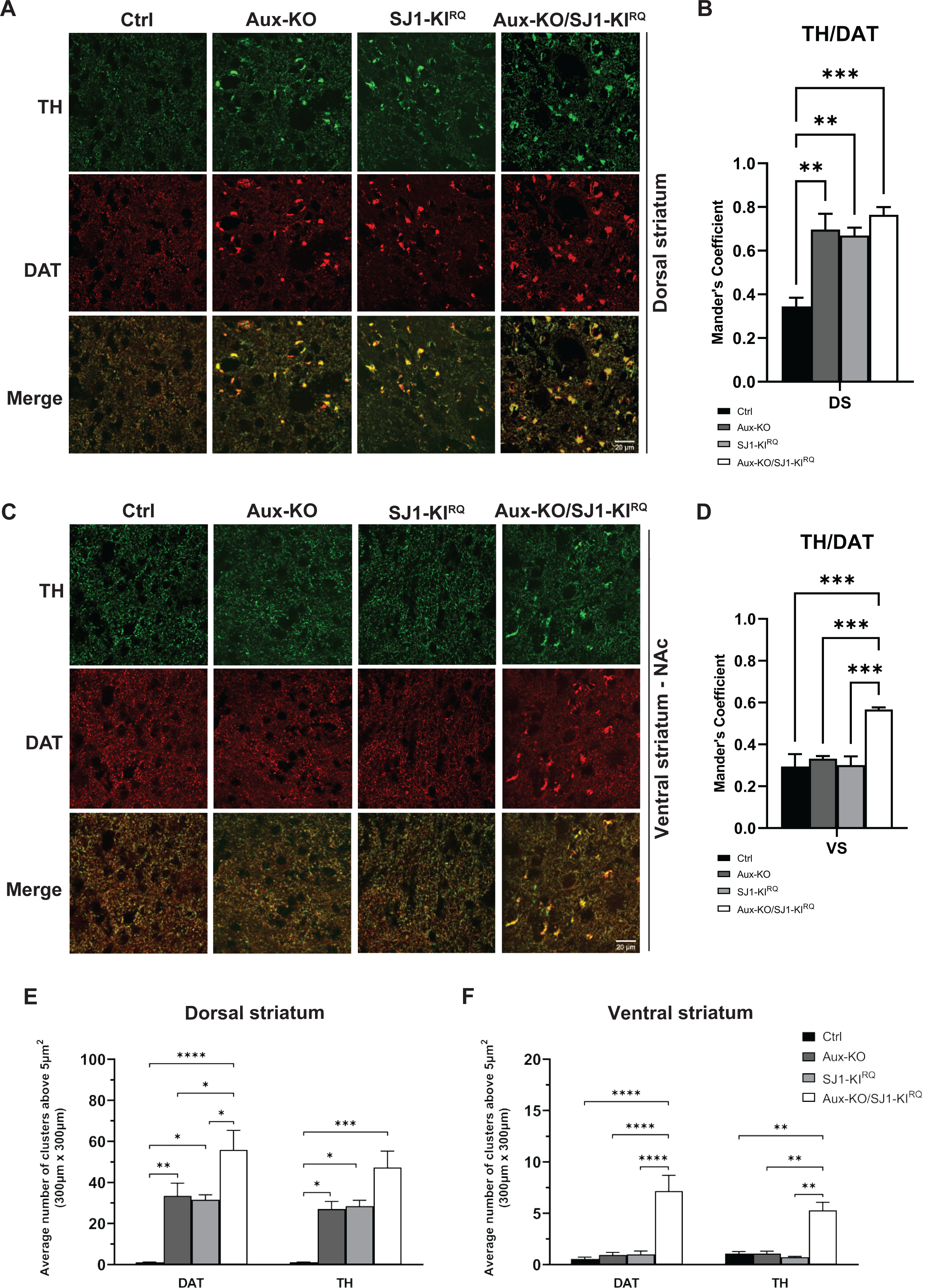
DAergic axon terminals undergo dystrophic changes in both dorsal and ventral striatum of Aux-KO/SJ1-KI^RQ^ mice. (A and C) Representative images for TH and DAT in cryopreserved-sections for 4 genotypes in (A) dorsal striatum and (C) nucleus accumbens (NAc) which is part of the ventral striatum. (B and D) Quantification of colocalization of TH and DAT clusters using Mander’s coefficient for (B) dorsal striatum, DS and (D) ventral striatum, VS. Data are represented as mean ± SEM (** *p*<0.01, *** *p*<0.001 by one-way ANOVA with post-hoc Tukey’s test). N= 3-4 mice per genotype. (E-F) Quantification for the average number of DAT and TH clusters above 5µm^2^ in (E) dorsal striatum and (F) ventral striatum of 1-2 months old mice. The number of clusters is calculated in five regions of interest (ROIs) which were selected at random for quantification of each mouse. Data are represented as mean ± SEM (* *p*<0.05, ** *p*<0.01, *** *p*<0.001, **** *p*<0.0001 by two-way ANOVA with post-hoc Tukey’s test). N = 3-4 mice for each genotype.

EM observations of SJ1-KI^RQ^ striata had shown that the presence of TH/DAT-positive clusters correlated with the occurrence of abnormal nerve terminals characterized by peculiar onion-like plasma membrane accumulations. Accordingly, similar structures were detected by EM in both auxilin KO and Aux-KO/SJ1-KI^RQ^ striata (Fig. 5). Moreover, EM analysis confirmed that the increase of TH/DAT-positive clusters corresponded to an increase in the number of onion-like plasma membrane derived structures. While these structures were difficult to be found in thin sections of single mutant mice due to their scattered distribution, they were much easier to be found in thin section of double mutant striata, confirming their abundance. Red arrowheads in Fig. 5 show that they represent plasma membrane invagination. Lamellae sometimes contained scattered small SV clusters or CCVs consistent with their axonal nature. Moreover, anti-TH immunogold labeling confirmed that these structures corresponded to TH accumulations (Supplementary Fig. 7).

**Figure 5.**
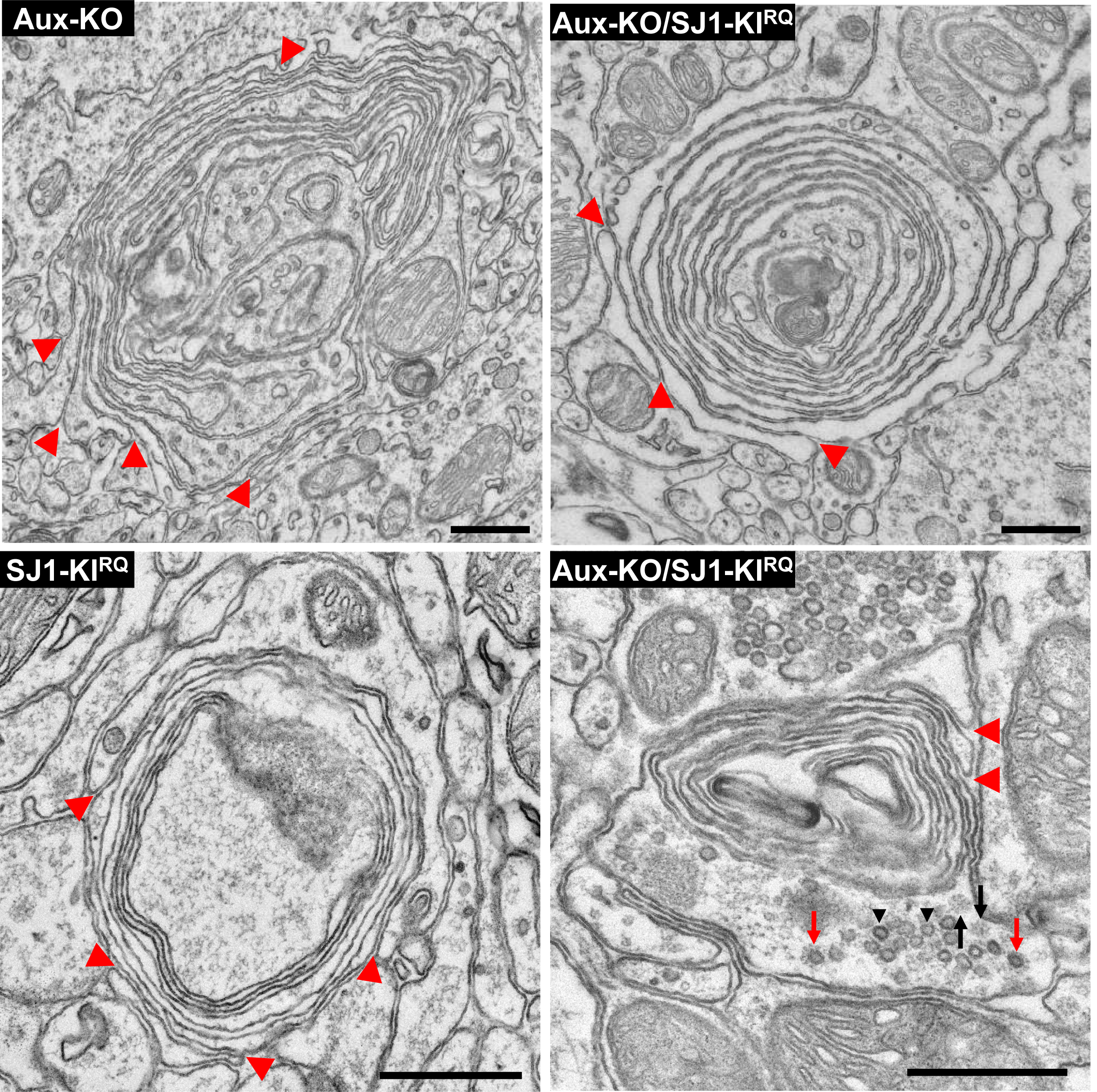
EM micrographs showing multilayered onion-like membrane structures in the dorsal striatum of Aux-KO, SJ1-KI^RQ^, and Aux-KO/SJ1-KI^RQ^ mice. These structures, which are positive for TH immunoreactivity (see Supplementary Fig 6), appear to result from invaginations of the plasma membrane as indicated by red arrowheads. Note in the bottom right field presence of a cluster of SVs (black arrowheads) and scattered CCVs (red arrows) and empty clathrin cages (black arrows), confirming that these structures represent dystrophic changes of nerve terminals. Scale bar: 500 nm.

Immunofluorescence for a variety of nerve terminal proteins and specific components of the DAergic system revealed co-enrichment of several other proteins with TH and DAT in these structures in Aux-KO/SJ1-KI^RQ^ mice. These include AADC (Fig. 6A, G), the plasma membrane marker SNAP25 (Fig. 6B, G) and a subset of SV proteins, such as synaptic vesicle glycoprotein 2C (SV2C) (Fig. 6C, G) and synaptotagmin 1 (Syt1) (Supplementary Fig. 8C) but not other SV house-keeping proteins such as SV2B (Fig. 6D, G) and synapsin (Supplementary Fig. 8D). Both AADC and SV2C were also clustered with TH/DAT in Aux-KO or SJ1-KI^RQ^ striatum (Supplementary Fig. 8A, B). Given the very low numbers of SVs in the onion-like structures but the abundance of plasma membrane, SV2C and Syt1 may have been more prone than others to become stranded in the plasma membrane in double mutant neurons. SV2C immunoreactivity was particularly strong consistent with a preferential expression of this SV2 isoform in the striatum and the midbrain, relative to SV2A and SV2B (http://dropviz.org/ and ^24, 25^). It is also worth to note that gene encoding SV2C was recently identified as a PD risk locus ^26, 27^. Importantly, the abnormal SNAP25 positive clusters were not observed in other brain regions, suggesting a selective impairment in the striatum (Supplementary Fig. 9A, B). None of the endocytic proteins tested, including clathrin light chain (CLC) and endophilin 1 were found in these aggregates (Fig. 6E, F, G), in agreement with the absence of accumulation of endocytic intermediates observed by EM at these sites.

**Figure 6.**
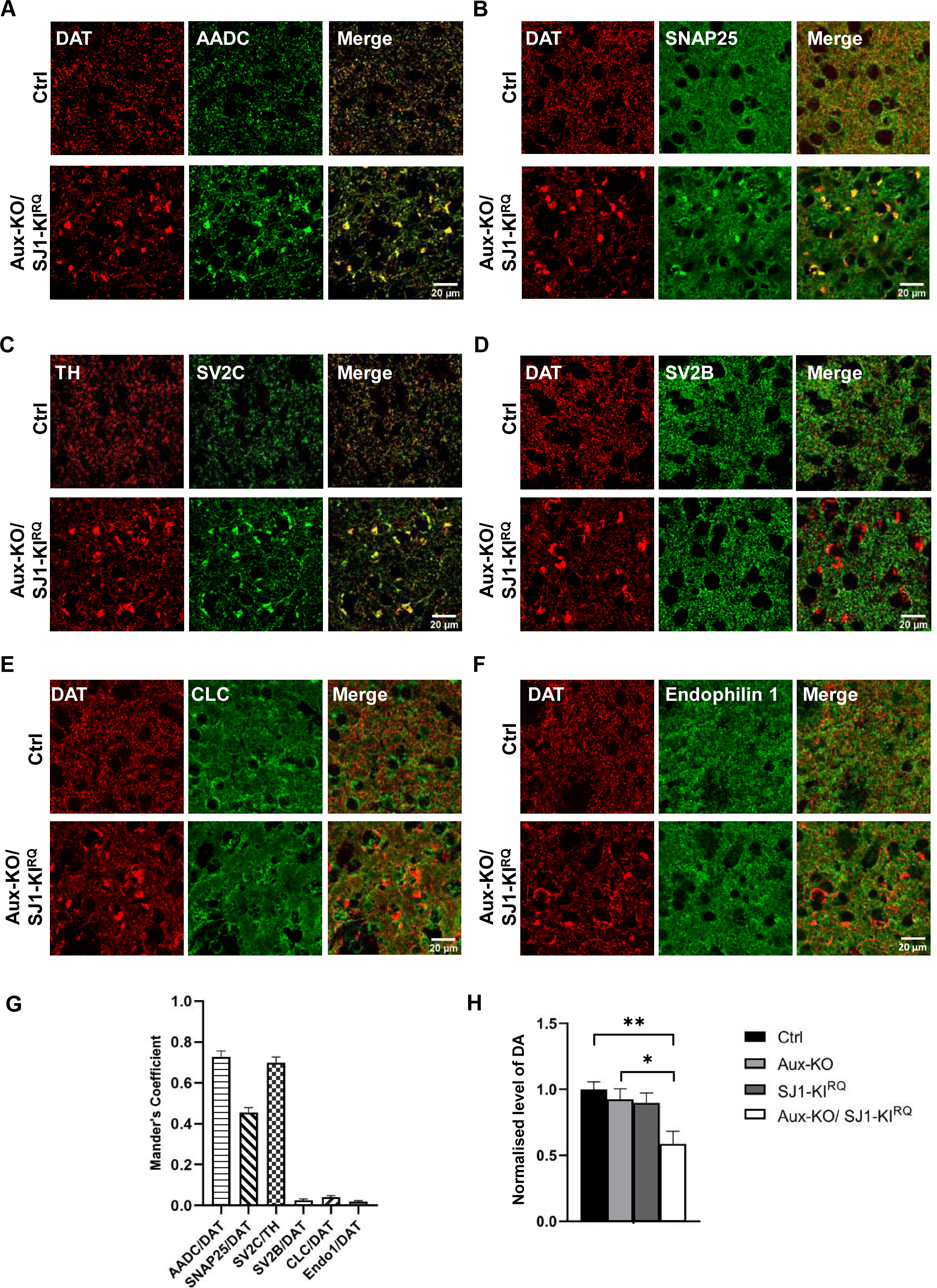
Accumulation of other proteins with TH/DAT in dystrophic DAergic nerve terminals in 1-month-old Aux-KO/SJ1-KI^RQ^ mice. (A) Double staining of DAT with AADC, a DA catabolism enzyme showed colocalization in the striatum of Aux-KO/SJ1-KI^RQ^ mice. (B) Double staining of DAT with SNAP25, a plasma membrane SNARE protein showed colocalization in the striatum of Aux-KO/SJ1-KI^RQ^ mice. (C-D) Double staining of SV2C (C) and SV2B (D) reveals that SV2C is specifically accumulated in TH/DAT-positive clusters in the striatum of Aux-KO/SJ1-KI^RQ^ mice, but not its family member SV2B. (E-F) Double staining of clathrin light chain (E) and endophilin (F) reveals that both endocytic proteins do not colocalize with TH/DAT-positive clusters in the striatum of Aux-KO/SJ1-KI^RQ^ mice. (G) Mander’s colocalization coefficient shows that AADC and SV2C colocalize the best with DAT/TH clusters, followed by SNAP25. SV2B, CLC and Endo1 do not colocalize with the DAT clusters. Data are represented as mean ± SEM. N = 5-11 random sampling sites. (H) Aux-KO/SJ1-KI^RQ^ shows 50% reduction in striatal DA levels measured using HPLC. Data are represented as mean ± SEM (* *p*<0.05, ** *p*<0.01 by one-way ANOVA with post-hoc Tukey’s test). N = 7 mice for each genotype.

Consistent with the occurrence of dystrophic changes of DAergic axons, measurements of the total content of DA in the striatum at 1-month old using HPLC, showed that DA levels were significantly decreased in striata of Aux-KO/SJ1-KI^RQ^ mice compared to those of single mutants and controls (Fig. 6H).

### Adaptive upregulation of striatal TH/AADC-positive interneuron and SV2C-positive cholinergic interneurons in Aux-KO/SJ1-KI^RQ^ mice

Most interestingly, we also detected some protein expression changes in local striatal neurons of Aux-KO/SJ1-KI^RQ^ mice. First, in sections of mutant striata immunolabeled with anti-TH antibody, in addition to the presence of TH-positive normal and abnormal nerve terminals which co-localized with DAT immunoreactivity, we also observed sparsely distributed TH-positive interneuron (THIN) cell bodies, different from Darpp32-positive medium spiny neurons (MSNs) which account for the predominant population in the striatum (Supplementary Fig. 10A). These neurons were not visible in WT and Aux-KO striata, and were only occasionally observed in SJ1-KI^RQ^ (Fig. 7A, B). Such neurons were positive for AADC (Fig. 7D, G), suggesting that they could possibly generate DA, however, they were also shown to be positive for GABA (Supplementary Fig. 10B), and negative for DAT (Fig. 7C, G). Second, we detected another neuronal population which was strongly labeled with two different SV2C antibodies in Aux-KO/SJ1-KI^RQ^ striata (Fig. 7E and Supplementary Fig. 10C, D). These SV2C-positive neuronal cell bodies did not overlap with TH/AADC-positive neurons, but were all positive for choline acetyltransferase (ChAT), a marker of striatal cholinergic interneurons (CHINs) (Fig. 7E, F, G and Supplementary Fig. 10C, D). Compared to WT sections, SV2C expression was much higher in a subset of ChAT-positive neurons in Aux-KO/SJ1-KI^RQ^ (Fig. 7H, I and Supplementary Fig. 10E, F). The intensity, number and distribution of ChAT remains unchanged between WT and Aux-KO/SJ1-KI^RQ^ (Fig. 7H, I), suggesting this is a specific upregulation of SV2C in the ChINs. We further examined other cell types in the striatum using immunofluorescence against Darpp32 for MSNs, GFAP for astrocytes and Iba1 for microglia. Both MSNs and glia cells appeared normal in morphology, localization and fluorescence intensity (Supplementary Fig. 8E, F, G). Collectively, these findings suggest that the local striatal microcircuitry has been affected in Aux-KO/SJ1-KI^RQ^ mutant mice, possibly as a result of DA deficiency and impaired DAergic innervation.

**Figure 7.**
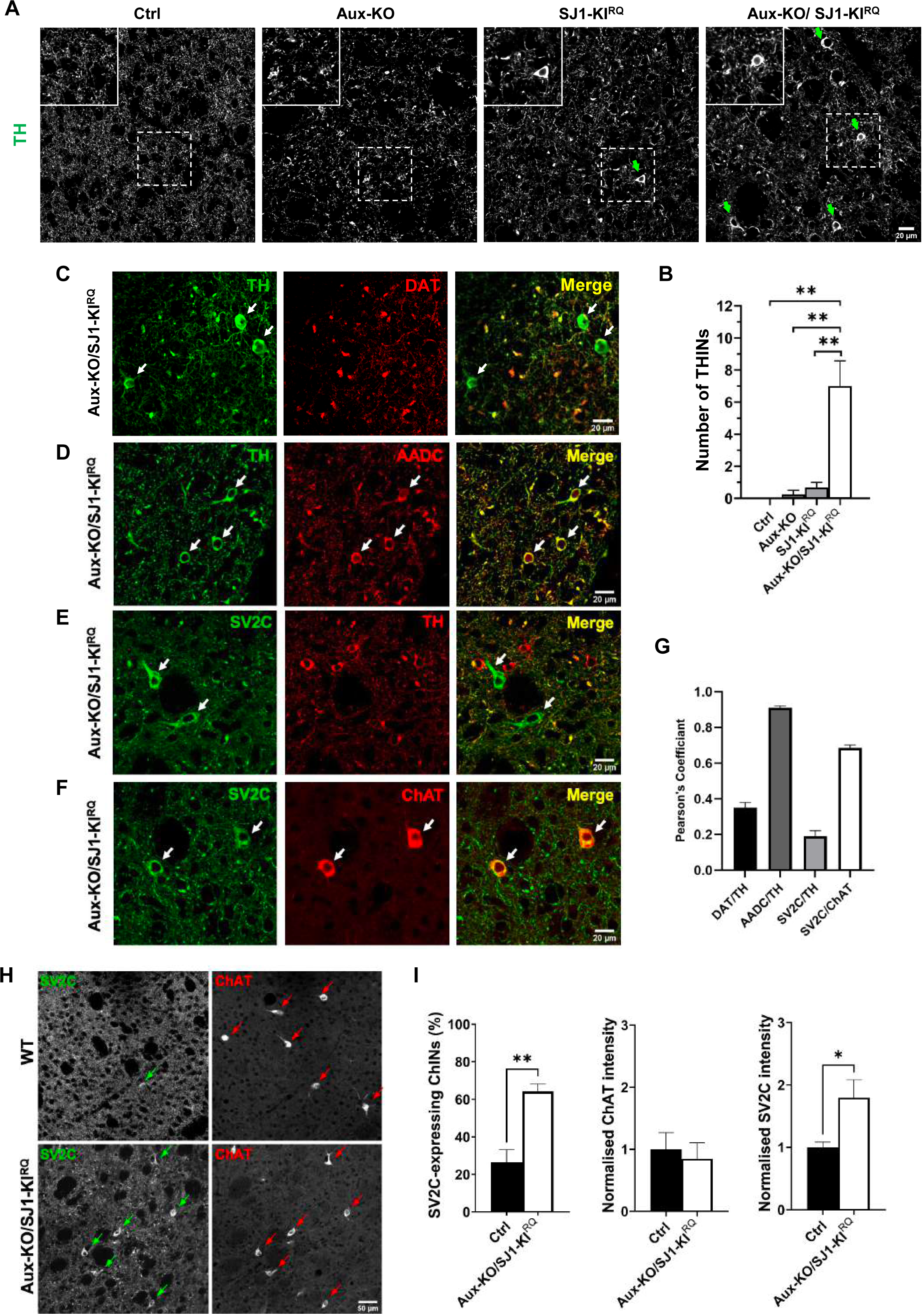
Adaptive changes of TH-positive interneurons and SV2C-positive cholinergic interneurons in Aux-KO/SJ1-KI^RQ^ striatum. (A) Immunostaining of TH reveals the Aux-KO/SJ1-KI^RQ^ striatum contains a large number of THINs. THINs are not found in Ctrl and Aux-KO striatum; and are less often found in SJ1-KI^RQ^ striatum. (B) Quantification for the number of THINs in 10 randomly selected regions of interest (ROIs) from a representative coronal section of the striatum. Data are represented as mean ± SEM (** *p*<0.01 by one-way ANOVA with post-hoc Tukey’s test). N = 3-4 mice for each genotype. (C) Immunostaining of TH (green) and DAT (red) reveals presence of TH-positive/DAT-negative neurons in the 1-month-old Aux-KO/SJ1-KI^RQ^ striatum. (D) THINs also show immunoreactivity towards anti-AADC antibody. (E) Representative images of SV2C and TH staining in the dorsal striatum of 1-month-old Aux-KO/SJ1-KI^RQ^ mice. TH and SV2C were observed to be expressed in separate neuron populations. SV2C-positive/TH-negative neurons are marked by white arrows. (F) Double staining of striatal region with cholinergic marker ChAT and SV2C shows overlap between the two in a specific neuronal subset, indicating that the SV2C-positive neurons are giant striatal ChINs. (G) The cell bodies of THINs and ChINs are selected as our region of interest to perform Pearson’s correlation coefficient. This demonstrate the two different types of interneurons: THINs where TH and AADC colocalizes well but not with DAT; ChINs where SV2C colocalizes well with ChAT but not with TH. Data are represented as mean ± SEM. N = 7-10 random sampling site. (H) Striatal IHC for single coronal section reveals subset of ChAT-positive ChINs that exhibit striking upregulation of SV2C in Aux-KO/SJ1-KI^RQ^ mice compared to control. (I) Quantification for percentage of ChAT-positive interneurons expressing SV2C, as well as individual intensity of SV2C and ChAT in striatal ChINs. For each individual mouse, 20 ChINs across a single coronal section of striatum were randomly selected for quantification. Data is represented as mean ± SEM (* *p*<0.05 and ** *p*<0.01, by Student’s unpaired *t* test). Ctrl: n=5 mice and Aux-KO/SJ1-KI^RQ^: n=5 mice.

## Discussion

In this study, we show that LOF mutations in two Parkinsonism-linked proteins implicated in endocytic clathrin-mediated budding at synapses, SJ1 and auxilin, produce similar dystrophic changes of nigrostriatal DAergic nerve terminals. Moreover, we show a strong synthetic interaction of SJ1 Parkinsonism mutation and auxilin KO. These findings provide striking evidence for the proposed functional partnership of the two proteins in clathrin coat shedding, with the phosphoinostide phosphatase activity of SJ1 being responsible for the dissociation of the clathrin adaptors from the membrane and auxilin playing a critical role, along with the chaperone HSC70, in the disassembly of the clathrin coat. In fact, the phosphoinositide phosphatase activity of SJ1 was proposed to help recruit auxilin. Importantly, they point to dysfunction in clathrin uncoating as one of the perturbations that can lead to juvenile/early-onset Parkinsonism. How such dysfunction in turn leads to disease remains to be further clarified. Mechanisms may include a defect in SVs recycling with an impact on DA release, other alterations of synaptic membrane traffic such as impairment of autophagosome formation and sequestration of critical factors by assembled clathrin. As SJ1 and auxilin are neuronal housekeeping proteins, a major open question is why is the DAergic system more severely affected, although, as previously shown by us and others, synaptic defects in SJ1-KI^RQ^ and auxilin mutant mice are not restricted to DAergic neurons.

Aux-KO mice have phenotypic manifestations that resemble those of SJ1-KI^RQ^ mice, and both of them recapitulate some of the manifestations of human Parkinsonism patients carrying mutations in the DNAJC6/PARK19 and the SYNJ1/PARK20 genes. However, the neurological defects of Aux-KO mice are less severe than those of SJ1-KI^RQ^ mice. This is probably due to the presence and compensatory upregulation of the auxilin paralogue auxilin-2/GAK in Aux-KO mouse brains ^19^. GAK itself was identified as a PD risk gene via a GWAS study ^28^.

Both the SJ1-KI^RQ^ mouse model and the Aux-KO model have been reported to show presynaptic clustering of endocytic factors and accumulations of assembled clathrin ^13, 19^. These clathrin structures were different in the two genotypes: primarily CCVs in SJ1-KI^RQ^ mice while empty clathrin cages in Aux-KO mice, as in the absence of auxilin spontaneously assembled cytosolic clathrin does not disassemble. These changes were widespread across the brain, but more prominent at inhibitory synapses, probably due to the tonic firing pattern in these synapses which requires more efficient SVs recycling, hence making these synapses more vulnerable to endocytic defects. A similar difference in the impact of endocytic protein mutations on excitatory and inhibitory synapses was also observed in SJ1 KO ^29^ and dynamin KO mice ^30^. The imbalanced excitatory/inhibitory synaptic transmission in SJ1-KI^RQ^ mice and Aux-KO mice could explain the epileptic seizures observed in both human patients and mice with mutations in SJ1 and auxilin.

The most interesting new observation that we have made in Aux-KO mice is the occurrence of histological defects in their striatum which are similar to those that we had observed in the striatum of SJ1-KI^RQ^ mice ^13^. These consist of TH/DAT-positive structures, also enriched in the plasma membrane protein SNAP25, that reflect presence of dystrophic DAergic nerve terminals and correlate with abnormal onion-like plasma membrane infoldings detectable by EM. The occurrence of these histological/cellular phenotypes in both mutant mouse models suggests a special liability of nigrostriatal DA neurons to perturbation of endocytic clathrin coat dynamics, which may be relevant to PD pathogenesis.

Possibly, as we discussed in our study of SJ1-KI^RQ^ mice, these onion-like invaginations of the plasma membrane are the result of endocytic impairment at a subset of synapses. However, they are not enriched with endocytic proteins nor several house-keeping SV proteins, consistent with the lack of enrichment of SVs in these terminals. Surprisingly, they are enriched in SV2C, which is the SV2 family member expressed in DAergic neurons and also a modulator of DA release ^24^. Interestingly, the gene encoding SV2C was recently identified by GWAS from both Asian and European cohorts as a PD risk locus ^26, 27^. Moreover, in both Aux-KO and SJ1-KI^RQ^ mice, dystrophic DAergic nerve terminals in the striatum peak in abundance at 1-2 months but gradually decreased (5 and 12 months), indicating either repair mechanisms or removal of degenerated nerve terminals in older mice. Interestingly, a follow up study on Parkinsonism patients carrying the SJ1 RQ mutation showed relative stability of clinical manifestations at later stages ^31^. The early-onset feature in both the Aux-KO and SJ1-KI^RQ^ mice also suggests there may be some neurodevelopmental defects in these mouse models. Indeed, Parkinsonism patients carrying both genes’ mutations suffer from developmental delay ^11, 12^, and a recent study of human midbrain-like organoids carrying the auxilin Parkinsonism mutation also showed impaired WNT-LMX1A signaling during DA neuron development ^21^.

The strong synthetic effect of the SJ1-KI^RQ^ mutation and the Aux-KO mutation is a key result that provides evidence for a similar pathogenetic mechanism resulting from mutations in these two proteins. Relative to single mutants, double mutant mice have much shorter lifespan and usually die between 3 to 4 weeks after birth, which is the critical period for neurodevelopment in rodents ^32^. Moreover, they display enhanced clustering of endocytic proteins and more abundant dystrophic TH/DAT/AADC/SV2C-positive nerve terminals, which additionally are not restricted to the dorsal striatum as in single mutant mice, but are also present in the ventral striatum. Thus, there appears to be a special vulnerability of DAergic neurons to the combined SJ1 and auxilin perturbations in the SN and VTA that project to the dorsal and ventral striatum, respectively. One factor that may contribute to the vulnerability of DAergic neuron to disruptions of two endocytic proteins may be their tonic firing activity, which may require efficient SVs recycling to sustain continuous release of DA. However, this is not a unique property of these neurons, suggesting that other factors come into play.

A striking finding in Aux-KO/SJ1-KI^RQ^ striata, besides the presence of dystrophic DAergic nerve terminals, was the occurrence of changes in two different types of interneurons, which likely reflect adaptive modifications. First, there was an increase in the number of TH-positive THINs. THINs were also positive for AADC, suggesting that they may synthesize DA. A similar increase was previously reported in neurotoxin-induced rodent and non-human primate PD models ^33, 34^, as well as in a genetic PD mouse model ^35^, and was interpreted as a compensatory change due to loss of DAergic terminals. The increase that we have observed in Aux-KO/SJ1-KI^RQ^ striata may have a similar explanation. However, DAT is not detected in these TH/AADC-positive interneurons, suggesting they are not “authentic” DA neurons ^36^. Second, in addition to the accumulation of SV2C with TH and DAT in DAergic terminals, we also observed upregulated SV2C expression in the soma of another subset of striatal interneurons, the ChAT-positive giant ChINs. Expression of SV2C in striatal ChINs has previously been reported ^37, 38^, but such expression is robustly increased in Aux-KO/SJ1-KI^RQ^ striata, while expression of ChAT remains relatively unchanged. Interestingly, a study involving neurotoxin PD mouse models to deplete DA also showed a significant increase in striatal SV2C mRNA levels ^39^. Considering that an imbalance of DAergic and cholinergic signals in the striatum is a feature of PD ^40^ and that SV2C has been linked by multiple studies to PD ^24, 26, 27, 39, 41^, this adaptive change in Aux-KO/SJ1-KI^RQ^ striata deserves further investigation.

In conclusion, we have reported here strong evidence for the hypothesis that mutations in SJ1 and auxilin may lead to Parkinsonism via a similar pathogenetic mechanism. Several PD causative genes have now been identified. The next goal is to identify the cellular process onto which these genes functionally converge, as such identification will provide insight into pathogenetic mechanisms with implication for therapeutic approaches. After the well-established partnership of PINK1 and Parkin, the cooperation of SJ1 and auxilin is the second clear example of functional partnership between two PARK genes. Additionally, we previously reported a synergistic function of SJ1 with another phosphoinositide phosphatase, Sac2, which GWAS studies had indicated as a candidate PD risk gene. Thus, we anticipate that further studies of auxilin and SJ1 will provide valuable insight into mechanisms that are at the core of at least some form of PD.

### Methods and Materials Animals

Both the Aux-KO (RRID: MMRRC_036980-JAX) and SJ1-KI^RQ^ mice were generated as described before ^13, 19^. These two mouse models were crossed with each other to generate the Aux-KO/SJ1-KI^RQ^ double-mutant mice. Mice were housed in SPF rooms with a 12-hour light/dark cycle. All experiments involving animals were conducted according to the protocol approved by SingHealth Institutional Animal Care and Use Committee (IACUC). WT or Heterozygous littermates of the mutant mice are used and defined as the control group.

### Antibodies

The following primary antibodies were obtained from Dr Pietro De Camilli’s lab at Yale University: rabbit anti-SJ1, rabbit anti-Auxilin, mouse anti-Amphiphysin 1, mouse anti-Clathrin Heavy Chain, rabbit anti-pan-Dynamin, rabbit anti-pan-Endophilin, mouse anti-GAD65, rabbit anti-SNAP25, rabbit anti-Synapsin, mouse anti-VAMP2, rabbit anti-Synaptophysin and mouse anti-Syt1. The other antibodies used in this study were obtained from commercial sources as stated: rabbit anti-LRRK2 (ab133474, RRID: AB_2713963) from Abcam; mouse anti-α-synuclein (610786, RRID: AB_2748880), mouse anti-AP2 (611350, RRID: AB_398872) and mouse anti-Hip1R (612118, RRID: AB_399489) from BD Biosciences; rabbit anti-DARPP-32 (2306, RRID: AB_823479) and rabbit anti-NPY (11976, RRID: AB_2716286) from Cell Signaling Technology; mouse anti-α-adaptin (MA1-064, RRID: AB_2258307) from Life Technologies; mouse anti-Amph2 (05-449, RRID: AB_309738), goat anti-ChAT (AB144P, RRID: AB_2079751), mouse anti-CLC (AB9884, RRID: AB_992745), rat anti-DAT (MAB369, RRID: AB_2190413), mouse anti-SV2C (MABN367, RRID: AB_2905667) and rabbit anti-TH (AB152, RRID: AB_390204) from Merck Millipore; mouse anti-β-actin (sc-47778, RRID: AB_2714189) and mouse anti-Hsc70 (sc-7298, RRID: AB_627761) from Santa Cruz Biotechnology; rabbit anti-Auxilin (HPA031182, RRID: AB_10611957), rabbit anti-GABA (A2052, RRID: AB_477652), rabbit anti-GFAP (ZRB2383, RRID: AB_2905668) and rabbit anti-SJ1 (HPA011916, RRID: AB_1857692) from Sigma-Aldrich; rabbit anti-Amphiphysin 1 (120002, RRID: AB_887690), rabbit anti-AADC (369003, RRID: AB_2620131), rabbit anti-SV2B (119102, RRID: AB_887803), rabbit anti-SV2C (119202, RRID: AB_887803), rabbit anti-Synaptogyrin 3 (103 302, RRID: AB_2619752), and rabbit anti-Syt11 (270003, RRID: AB_2619994) from Synaptic Systems; rabbit anti-Iba1 (019-19741, RRID: AB_839504) from FUJIFILM Wako Chemicals.

Secondary antibodies used were all purchased from commercial sources as stated: donkey anti-mouse IgG (H+L) Alexa Fluor 594 (A21203, RRID: AB_141633), goat anti-mouse IgG (H+L) Alexa Fluor 488 (A11001, RRID: AB_2534069), goat anti-mouse IgG (H+L) Alexa Fluor 594 (A11032, RRID: AB_2534091), goat anti-mouse IgG (H+L) Alexa Fluor 647 (A21236, RRID: AB_2535805), donkey anti-rabbit IgG (H+L) Alexa Fluor 488 (A21206, RRID: AB_2535792), goat anti-rabbit IgG (H+L) Alexa Fluor 488 (A11034, RRID: AB_2576217), goat anti-rabbit IgG (H+L) Alexa Fluor 594 (A11037, RRID: AB_2534095), goat anti-rabbit IgG (H+L) Alexa Fluor 647 (A21244, RRID: AB_2535812), goat anti-rat IgG (H+L) Alexa Fluor 488 (A11006, RRID: AB_2534074), goat anti-rat IgG (H+L) Alexa Fluor 594 (A11007, RRID: AB_10561522) and donkey anti-goat IgG (H+L) Alexa Fluor 488 (A11055, RRID: AB_2534102) from Life Technologies and IRDye 800CW donkey anti-rabbit IgG (926-32213, RRID: AB_621848), IRDye 800CW donkey anti-mouse IgG (926-32212, RRID: AB_621847), IRDye 680RD donkey anti-mouse IgG (926-68072, RRID: AB_10953628) and IRDye 800CW goat anti-rat (926-32219, RRID: AB_1850025) from LI-COR Biosciences.

### Motor Behavioral tests

Aux-KO and littermate control mice between 2-13 months old were used for balance beam and rotarod tests. The mice were divided into three age groups: 2-4 months old (n=13-15), 5-8 months old (n=7-12) and more than 9 months old (n=8-15) for each genotype. Both males and females were used for the behavioral assays. Both tests were conducted during the light period. The mice were allowed to acclimatize to the test for at least 30 min before each test.

#### Balance Beam Test

A narrow beam (10 mm width) was suspended 15 cm above a soft padding. Mice were placed on end of the beam and during their trip towards the other end, the number of missteps (paw slips) was recorded. The test was conducted thrice for each mouse and the average number of missteps was calculated.

#### Rotarod Test

Mice were placed on a rod which is rotating at 4 rpm with an acceleration to 40 rpm within 6 min. The duration of the mice staying on the rod is measured two times at 10 min intervals and the average ‘latency to fall’ was determined for each mouse.

### Immunoblotting

Mouse brain tissues were homogenized in buffer containing 20 mM Tris-HCl (pH 7.4), 150 mM NaCl, 2 mM EDTA and cOmplete^TM^ EDTA-free Protease Inhibitor Cocktail (Roche, USA). The homogenized samples were centrifuged at 700 g for 10 min to obtain the post-nuclear supernatant (PNS). Protein concentration was determined by the Pierce BCA Protein Assay Kit. SDS-PAGE and western blotting were performed by standard procedures. The membranes were blotted with primary antibodies and then IRDye® Secondary Antibodies (LI-COR, USA). The protein bands were detected using Odyssey ® CLx imaging system and its intensity was quantified via Image Studio Lite software (Ver 5.2, RRID: SCR_013715). A minimum of three independent sets of samples were used for quantification.

### Primary Cortical Neuron Culture and Staining

Cultures of cortical neurons were prepared from P0 to P2 neonatal mouse brains by previously described methods ^42^ and used at DIV19. Cortical tissue was dissected out, placed in ice-cold HBSS, and diced into small pieces of less than 1 mm^3^ per piece. Tissue was then digested for 30 min in an activated enzyme solution containing papain (20 U/ml) and DNase (20 μg/ml) at 37°C, followed by gentle trituration. The cell suspension is centrifuged at 300 g for 5 min. The cell pellet is resuspended and plated onto poly-d-lysine-coated coverslips to a density of 60,000 cells/cm^2^. 2-3 hours after plating, the medium was exchanged to Neurobasal/B27 serum-free medium, and cells were maintained at 37°C in a 5% CO2 humidified incubator. Cells were fixed with 4% paraformaldehyde (PFA) and 4% sucrose in 1 X PBS for 20 min, followed by 10 min incubation with 50 mM NH_4_Cl, then blocked and permeabilized with 5% bovine serum albumin (BSA), 1 X phosphate-buffered saline (PBS) and 0.1% Triton X-100. Primary and secondary antibody incubations were subsequently performed in the same buffer. Alexa 488, 594 and 647 conjugated secondary antibodies were purchased from Invitrogen. After washing, samples were mounted on slides with Fluoromount-G (Invitrogen). Samples were observed and imaged on a spinning disk system (Gataca Systems) based on an inverted microscope (Nikon Ti2-E; Nikon) equipped with a confocal spinning head (CSU-W, Yokogawa) and a Plan-Apo 60x oil TIRF objective. The same exposure time and laser intensity were used when imaging the same marker for all 4 genotypes.

### Brain Histology and Immunofluorescence

Mice were anesthetized with a Ketamine/Xylazine anesthetic cocktail injection, perfused transcardially with ice-cold 4% PFA in 1X PBS and the brains were kept in the same fixative overnight at 4°C. On the next day, brains were transferred to 30% sucrose in 1 X PBS and kept overnight at 4°C on top of a roller. Brains were then embedded in OCT (Tissue-Tek) and freeze in liquid nitrogen-cooled 2-methylbutane (Isopentane). Coronal or sagittal sections of 20 µm thickness were cut with a cryostat. The sections are either mounted on adhesive slide, SuperFrost® Plus (VWR) or collected as floating sections in 1 X PBS in a 24-well plate. Sections were then blocked in 5% BSA and 0.1% Triton X-100 in 1 X PBS for 1 hour at room temperature. After blocking, sections were incubated with primary antibodies (diluted in the same buffer) and kept overnight at 4°C. Subsequently, sections were washed 3 times for 10 min with 0.1% Triton X-100 in 1 X PBS, then incubated with Alexa-conjugated secondary antibodies for 1 hour at room temperature. Finally, the sections were mounted with Fluoromount-G with or without DAPI (Invitrogen) and sealed with nail polish. The floating sections were mounted onto slides after all the staining procedures. Images were acquired with a spinning disk system (Gataca Systems) based on an inverted microscope (Nikon Ti2-E; Nikon) equipped with a confocal spinning head (CSU-W, Yokogawa) and a Plan-Apo 40x oil objective. The same exposure time and laser intensity were used when imaging the same marker for all 4 genotypes.

For stereological analysis of TH-positive dopaminergic neurons in the midbrain, 30 µm coronal section were collected, incubated with 0.1% H_2_O_2_ for 20 minutes to quench endogenous peroxidase activity and subsequently incubated with buffer containing 5% BSA, 1X PBS and 0.1% Triton X-100 for 1 hour at room temperature. Next, sections were incubated overnight at 4°C with anti-TH primary antibody (1:1000). The sections were then washed with 1X PBS 3 times for 10 minutes and incubated with a biotinylated anti-rabbit secondary antibody (Vector Laboratories, USA) for 1 hour at room temperature, later followed by incubation with Avidin/Biotin complex (ABC) reagent for 45 minutes at room temperature (Vector Laboratories PK-6200). Finally, immunoreactivity was revealed by incubation with diaminobenzidine (DAB) (Vector Laboratories SK-4100). Stereological analysis was performed using the optical fractionator probe in the Stereo Investigator software (MBF Bioscience, USA). A total of 9 coronal sections (every 4^th^ serial section across the midbrain) were collected for counting. The substantia nigra par compacta (SN) and ventral tegmental area (VTA) regions were outlined based on the Allen mouse brain atlas using 5x objective lens and counts were performed using 60x oil objective lens. The parameters used include a counting frame size of 50 x 50 μm, a sampling site of 132 x 71 μm, a dissector height of 13 μm, 2 μm guard zones and coefficient of error (Gunderson, m=1) were less than 0.1.

### Quantification of Immunoreactivity Clustering and TH-positive Striatal Interneurons

The endocytic protein clustering quantification was conducted on the Fiji software (Version ImageJ 1.52p/Java 1.8.0_172 (64-bit), RRID: SCR_002285) as follows. The same threshold intensity was applied to all images and a random region of interest with an area of 500 μm^2^ was manually selected. A mask was used to selectively quantify puncta larger than 1 μm^2^. The Fiji plugin “Analyze particles” was then used to measure average fluorescence intensity of the puncta.

The quantification of TH/DAT-positive clustering in striatum was also done similarly using Fiji software. In this case, for each mouse, 5 random sampling sites were selected respectively for both dorsal and ventral striatum. At least three mice from each genotype and each age group were collected for this purpose. The threshold setting for the clusters was set to quantify puncta bigger than 5 μm^2^ to ensure normal positive axonal terminals were not counted in. The average number of clusters was counted automatically with Fiji plugin “Analyse Particles” based on all five random sampling sites for each mouse.

Quantification of the THINs were done with the same images used to quantify TH/DAT-positive clusters. The number of TH-positive cell bodies were counted from the 5 random sampling sites selected for the dorsal and ventral striatum respectively.

### Quantification of Striatal Cholinergic Interneurons (ChINs)

The number of ChINs was quantified from individual coronal sections of the striatum double stained with ChAT and SV2C. Image acquisition involved tiling of the entire section using a 20x Air lens, keeping the same laser intensity and exposure for both channels on MetaMorph Microscopy Automation and Image Analysis Software (RRID: SCR_002368). Stitched images were then set to the same threshold intensity for each individual channel in Fiji software (Version ImageJ 1.52p/Java 1.8.0_172 (64-bit), RRID: SCR_002285). Cell Counter plugin was used to count visible cell bodies across the striatum (outlined on the basis of Allen mouse brain atlas). Percentage ratio of the cell count from individual channels was used as a measure of ChINs expressing SV2C.

Quantification of ChAT and SV2C intensities in ChINs was done with the same images used for quantifying number of cholinergic interneurons. Manually drawn ROIs were used to analyse the intensities of 20 neurons randomly selected across the striatum. Corrected total cell fluorescence (CTCF) was calculated and the intensities of each channel were normalised to averaged control levels. For both genotypes, five individual mice were analysed for this purpose.

### Colocalization analysis

Colocalization analysis was also performed on the Fiji software (Version ImageJ 1.52p/Java 1.8.0_172 (64-bit), RRID: SCR_002285). For the quantification of colocalization of immunofluorescence signals in the clusters (both cortical endocytic clusters and striatal DAergic TH/DAT-positive clusters), the images are set to the same threshold to mask all the normal, healthy axon terminals. For colocalization analysis of Amph1/Gad65, 8-10 neurons were quantified for each genotype. For colocalization analysis of TH/DAT, 5 random sampling sites were selected respectively for both dorsal and ventral striatum of each mouse. At least three mice from each genotype were collected for this purpose. For the colocalization analysis of other markers with TH or DAT, 5-11 random sampling sites were selected from the dorsal striatum. Fiji plugin, JACoP ^43^, was used to set the threshold and calculate Mander’s coefficient. Only M1 values are reported. For the colocalization of immunofluorescence cell bodies’ signal of THINs and ChINs, 7-10 soma from 3-4 random regions of interest (ROIs) of striatum were selected. JACoP was used to calculate Pearson’s coefficient for colocalization analysis.

### Electron Microscopy (EM)

All EM reagents were from EMS, Hatfield, PA, unless noted otherwise. 1-3 months old mice were anesthetized and fixed by transcardial perfusion with 4% formaldehyde and 0.125% glutaraldehyde in 0.1 M PB buffer. Brain were removed and dissected in small pieces (0.5 × 0.5 x 0.5 mm^3^) and further incubated in the same fixative at 4 °C overnight. Brain tissues were further incubated in 2.5% glutaraldehyde in 0.1 M sodium cacodylate buffer for additional 1 hour at room temperature, post-fixed (1 hour) in 2% OsO_4_, 1.5% K_4_Fe(CN)_6_ (Sigma-Aldrich, St. Louis, MO) in 0.1M sodium cacodylate buffer, stained (overnight) with 2% aqueous uranyl acetate at 4 °C, dehydrated in gradually increasing concentration of EtoH, and embedded in Embed 812. Ultrathin sections, about 60 nm thick, were cut with a Leica ultramicrotome and examined in a Talos L120C TEM microscope at 80 kV. Images were taken with Velox software and a 4k × 4K Ceta CMOS Camera (Thermo Fisher Scientific). For quantification, images of synapses were selected based on the presence of an active zone. The number of SVs (diameter ≤ 80nm), CCVs, and clathrin cages was measured in more than 120 synapses for each control and mutant condition. These values were normalized to the cross-sectional area of the presynaptic terminal. Results of the morphometric analysis are presented as mean ± SEM. Statistical significance was evaluated using one-way ANOVA followed by Games-Howell’s multiple comparison test.

### Immunogold labelling

Aux-KO/SJ1-KI^RQ^ mice were transcardially perfused with 4% formaldehyde and 0.125% glutaraldehyde in 0.1M phosphate buffer (pH 7.4). Dorsal striata were dissected and embedded in 1% gelatin in 0.1M phosphate buffer. Tissue pieces were trimmed, infiltrated in 2.3M sucrose, and then were frozen rapidly onto aluminium pins in liquid nitrogen. 60nm frozen sections on carbon/formvar coated grids were prepared with a Leica Cryo-EMUC6 UltraCut microtome. Sections were labelled with rabbit anti-Tyrosine Hydroxylase (TH) and 10nm Protein A gold (Utrecht Medical Center) ^44^. Grids were examined in FEI Tecnai Biotwin TEM at 80Kv. Images were taken with Morada CCD and iTEM (Olympus) software.

### HPLC analysis to detect striatal levels of dopamine

Striatum tissues dissected from mice brain were homogenized in 0.5N perchloric acid with 100µM of deferoxamine mesylate and 100µM of glutathione. The homogenized samples were further sonicated, centrifuged and the supernatants were filtered using 0.1 µm PVDF centrifugal filters before collecting the filtrates for HPLC analysis. A reversed-phase UltiMate 3000 HPLC system (Thermo Fisher Scientific) with an electrochemical detector and a reversed-phase column (Vydac Denali, C18, 4.6 x 250mm, 5µm particle size, 120 Å pore size) were used to run the samples. The HPLC run was performed at a flow rate of 0.5 ml per minute with a mobile phase containing 1.3% sodium acetate, 0.01% EDTA (pH8.0), 0.5% sodium 1-heptanesulfonate, 7% acetonitrile (v/v) and 2% methanol (v/v), adjusted to pH 4.0 with acetic acid. All buffers used for HPLC analysis were double filtered through 0.2 µM nylon membranes. Dopamine in the samples was identified by retention time of dopamine standard (around 14 min) and quantified by measuring the area under the peak using the software Chromeleon^TM^ 7.2 Chromatography Data System (Thermo Fisher Scientific). The areas under the peaks were normalised to their respective tissue weight. 7 mice were used for each genotype for quantification.

### Statistical Analysis

All statistical analysis was performed using GraphPad Prism (Ver 9.3.1, RRID: SCR_002798). All graphs were also plotted on GraphPad Prism. Data are presented as mean ± SEM unless otherwise stated. Statistical significance was determined using the Student’s unpaired *t* test for the comparison of two independent groups or ANOVA with Tukey’s Honest Significant Difference or Games-Howell’s post hoc test for multiple group comparisons. Data with *p*-values <0.05, <0.01, <0.001, and <0.0001 are represented by asterisks *, **, ***, and ****.

## ABBREVIATIONS

AADC: Aromatic l-amino acid decarboxylase
AP2: Adaptor protein complex 2
CCVs: Clathrin coated vesicles
ChAT: Choline acetyltransferase
ChINs: Cholinergic interneurons
DA: Dopamine
DAergic: Dopaminergic
DAT: Dopamine transporter
GAK: Cyclin G-associated kinase
GFAP: Glial fibrillary acidic protein
GWAS: Genome-wide association study
H&E: Hematoxylin and eosin
Iba1: Ionized calcium-binding adapter molecule 1
KI: Knock-in
KO: Knockout
LOF: Loss-of-function
MSNs: Medium spiny neurons
NPY: Neuropeptide Y
PD: Parkinson’s Disease
SJ1: Synaptojanin 1
SN: Substantia nigra
SVs: Synaptic vesicles
SV2: Synaptic vesicle glycoprotein 2
Syt1: Synaptotagmin 1
THINs: TH-positive interneurons
VTA: Ventral tegmental area

## Acknowledgment

We thank Morven Graham for technical help and Prof. Zhang Su-Chun for sharing the reagents. This research was supported in part by grants from Ministry of Education Academic Research Fund (AcRF) Tier 2 Singapore (MOE2019-T2-2-090) to MC, from the NIH (NS36251 and DA18343), the Kavli Institute for Neuroscience, the Parkinson Foundation and the Aligning Science Across Parkinson’s grant ASAP-000580 through the Michael J. Fox Foundation for Parkinson’s Research (MJFF) to PDC. For the purpose of open access, the author has applied a CC BY public copyright license to all Author Accepted Manuscripts arising from this submission.

## Author Contributions

X.Y.N., P.D.C. and M.C. designed the experiments. X.Y.N., Y.W., Y.L., S.M.Y and M.C. performed the experiments. X.Y.N., P.D.C. and M.C. wrote the paper. L.E.G. provided Aux-KO mice. All authors read and approved the final manuscript.

## Availability of data and materials

The datasets generated and/or analyzed in this study are available from the corresponding author Mian Cao on request.

## Supplementary Figure Legends

**Supplementary Figure 1. Aux-KO/SJ1-KI^RQ^ mice are smaller in size and has more severe DAergic phenotype at 7-month-old.**

(A-B) Image of the size of P25 Aux-KO/SJ1-KI^RQ^ mice comparing with its littermate single mutants; (A) Aux-KO and (B) SJ1-KI^RQ^.

(C) Immunostaining of NPY shows elevated expression of NPY in the dentate gyrus of Aux-KO/SJ1-KI^RQ^ mice.

(D) Quantification of fluorescence intensity of NPY staining shown in (C). Data are represented as mean ± SEM (* *p*<0.05 by one-way ANOVA with post-hoc Tukey’s test). N = 3 mice for each genotype.

(E) Representative images for double immunofluorescence of TH/DAT, AADC/DAT and SV2C/DAT for all 4 genotypes at 7-month-old. The only Aux-KO/SJ1-KI^RQ^ that managed to survive till this age showed increased number of clusters, THINs and ChINs compared to the other 3 genotypes.

**Supplementary Figure 2. Aux-KO/SJ1-KI^RQ^ mice have normal brain development and architecture.**

H&E staining reveals absence of gross structural abnormality in various regions (cortex, hippocampus, striatum and cerebellum) of 1-month-old SJ1-KI^RQ^, Aux-KO and Aux-KO/SJ1-KI^RQ^ mice brain.

**Supplementary Figure 3. Absence of gliosis in Aux-KO/SJ1-KI^RQ^ brain.**

(A and B) No obvious difference in immunostaining intensity of (A) Iba1-microglia marker and

(B) GFAP-astrocytes marker in different regions of 1-month-old Ctrl, SJ1-KI^RQ^, Aux-KO and Aux-KO/SJ1-KI^RQ^ mice brain.

**Supplementary Figure 4. Western blot analysis of endocytic proteins, synaptic proteins, DAergic markers and PD-related proteins in mouse brains.**

(A) Representative blots for various proteins involved in PD and synaptic endocytosis from 1-month-old control, SJ1-KI^RQ^, Aux-KO and Aux-KO/SJ1-KI^RQ^ whole brain homogenates.

(B) Quantification of expression levels of the proteins shown in (A). Protein levels were normalized to the level of Actin. Data are represented as mean ± SEM (one-way ANOVA with post-hoc Tukey’s test). N = 3-4 mice for each genotype.

**Supplementary Figure 5. Presence of dystrophic changes only in the axon terminals of DAergic neurons but not the cell bodies of Aux-KO/SJ1-KI^RQ^ mice.**

(A) Double staining of anti-TH and anti-AADC showed normal morphology of DAergic neuron cell bodies and dendrites in the midbrain.

(B) Stereological analysis of TH-positive cell bodies in the SN and VTA region of control and Aux-KO/SJ1-KI^RQ^ mice at 1 month old. The estimated cell numbers in one hemisphere is shown. Data are represented as mean ± SEM (Student’s unpaired t-test). n=7 mice for control whereas n=6 mice for Aux-KO/SJ1-KI^RQ^.

(C) AADC (green) and DAT (red) immunostaining reveals the presence of AADC/DAT-positive aggregates in the olfactory tubercle (OT, part of ventral striatum) of Aux-KO/SJ1-KI^RQ^ mice.

**Supplementary figure 6. Overview of cluster distribution in the striatum of Aux-KO, SJ1-KI^RQ^ and Aux-KO/SJ1-KI^RQ^ mice.**

Tiling of a single coronal striatum section immunostained for DAT. White dotted line separates the dorsal and ventral striatum of the mice. Insets on the bottom left corner of the image showed high magnification images of the clusters for dorsal (pink, top) and ventral (purple, bottom) striatum for each genotype. Note the absence of clusters in the ventral striatum in all mice except the Aux-KO/SJ1-KI^RQ^ mice. Bottom right inset of the Aux-KO tiling image depicts the dorsal (pink) and ventral (purple) mouse striatum.

**Supplementary figure 7. EM immunogold labelling of onion-like membrane structure in dorsal striatum of Aux-KO/SJ1-KI^RQ^ mice.**

(A) Anti-TH immunogold labelling (10 nm gold particles) of an ultrathin frozen sections of Aux-KO/SJ1-KI^RQ^ dorsal striatum. The multilayered membrane structures are positive for TH immunoreactivity (boxed in white box). Scale bar: 1 µm

(B) The onion-like membrane structures boxed in white in (A) is enlarged and shown here. Scale bar: 250 nm

**Supplementary figure 8. Various other markers examined with TH/DAT-positive clusters.**

(A-B) Representative images for immunoreactivity of anti-DAT or TH with (A) anti-AADC and

(B) anti-SV2C in Aux-KO and SJ1-KI^RQ^ mice.

(C) Immunostaining of DAT with Syt1 showed partial colocalization in Aux-KO/SJ1-KI^RQ^ double mutant striatum.

(D-G) Double immunofluorescence of synapsin, GFAP, Iba1 and Darpp32 with TH/DAT showed no colocalization in the striatum of Aux-KO/SJ1-KI^RQ^ mice.

**Supplementary figure 9. Immunofluorescence analysis of SNAP25 in different brain regions.**

(A) Immunostaining of SNAP25 (green) showed partial colocalization with DAT-positve clusters (red) in the striatum of Aux-KO, SJ1-KI^RQ^ and Aux-KO/SJ1-KI^RQ^ mice.

(B) Immunoreactivity of anti-SNAP25 in the cortex of WT, Aux-KO, SJ1-KI^RQ^ and Aux-KO/SJ1-KI^RQ^ mice. No clusters of SNAP25 were observed in the cortex.

**Supplementary figure 10. Properties of striatal THINs and ChINs in the striatum of Aux-KO/SJ1-KI^RQ^ mice.**

(A) Immunoreactivity of anti-TH with anti-Darpp32 showed that THINs is a distinct group of neurons from Darpp32-positive MSNs.

(B) Double staining of anti-TH and anti-GABA showed that THINs are GABAergic striatal interneurons.

(C-D) Immunostaining of SV2C using a different anti-SV2C antibody (host species: Rabbit) labelled the same structures: (c) ChAT-positive ChINs and (d) SV2C/DAT-positive clusters in Aux-KO/SJ1-KI^RQ^.

(E-F) A large microscope field of view showed the distribution of (e) SV2C-positive and (f) ChAT-positive interneurons in the striatum of WT and Aux-KO/SJ1-KI^RQ^ mice.

## Supplementary Video Legends

**Supplementary Video 1 Tonic-clonic epileptic seizures in a 5-month-old Aux-KO mouse.**

**Supplementary Video 2 Balance beam test in 2-month-old (A) Ctrl and (B) Aux-KO mice.**

**Supplementary Video 3 Balance beam test in 4-month-old (A) Ctrl, (B) Aux-KO/SJ1-WT, (C) Aux-Ht/SJ1-KI^RQ^ and (D) Aux-KO/SJ1-KI^RQ^ mice.** This 4-month-old Aux-KO/SJ1-KI^RQ^ mouse is the same mouse that managed to survive till 7-month-old.

