## Supplementary figures for "Mutations in Parkinsonism-linked endocytic proteins synaptojanin1 and auxilin have synergistic effects on dopaminergic axonal pathology"

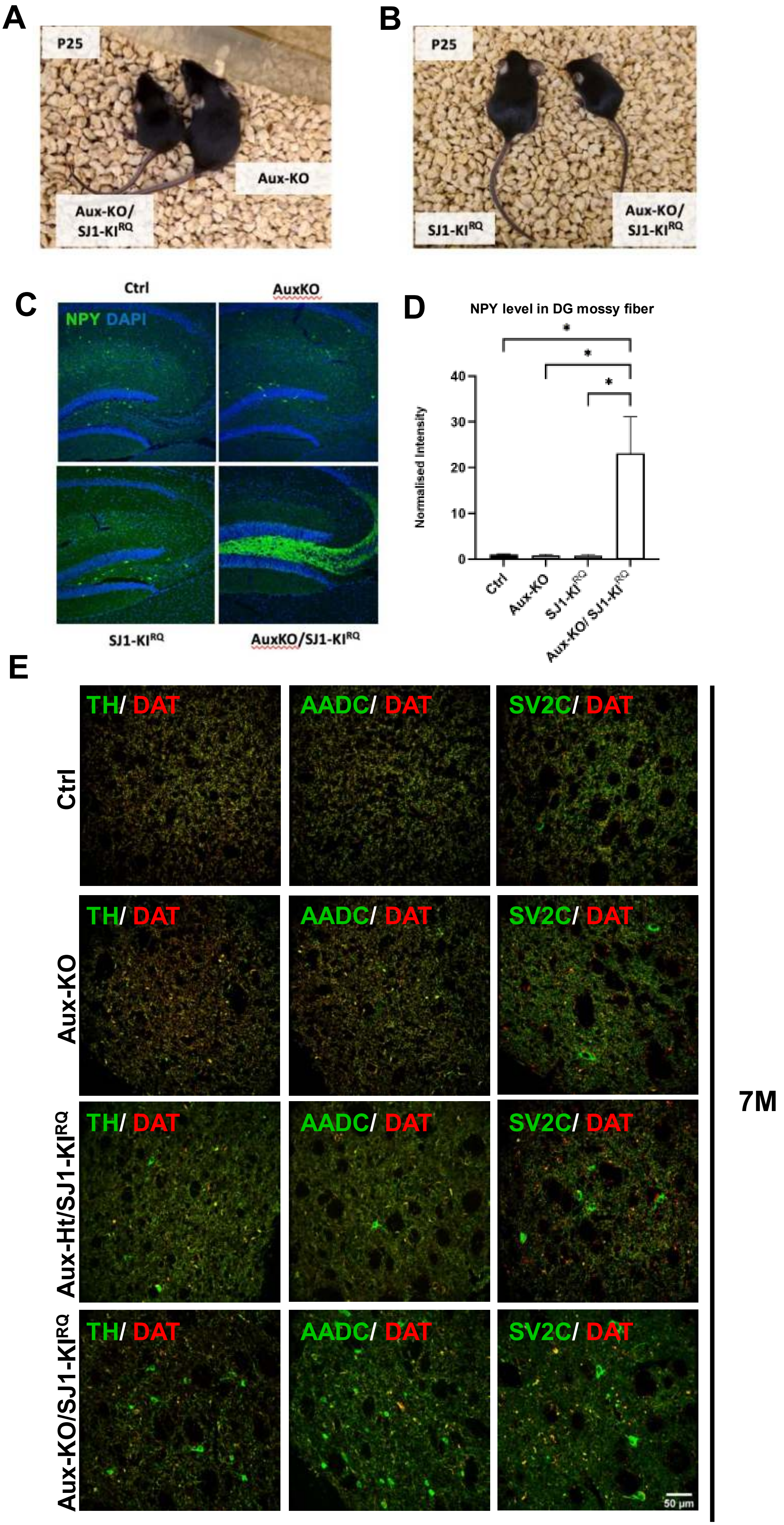


**Supplementary Fig 1**

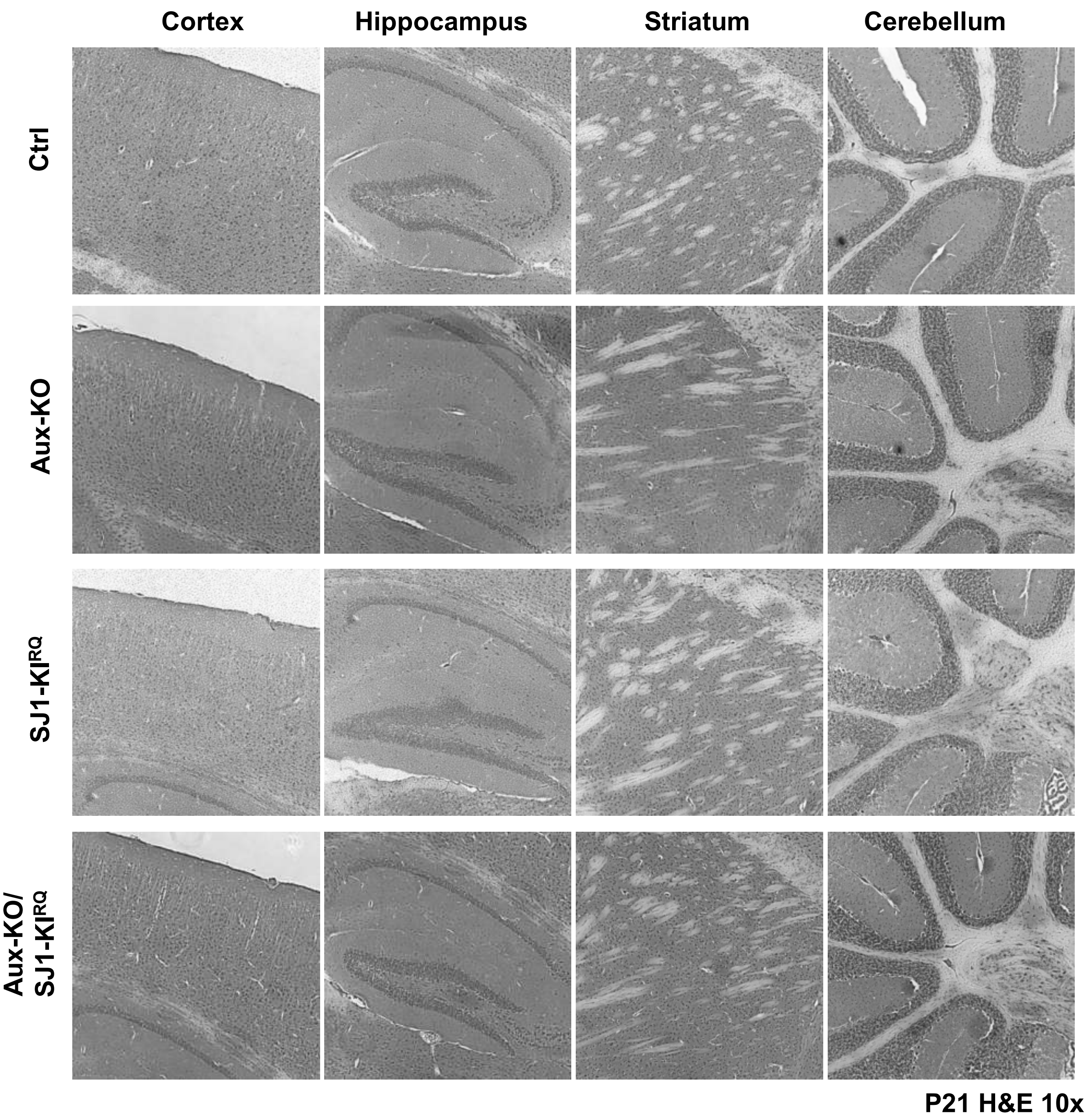


**Supplementary Fig 2**

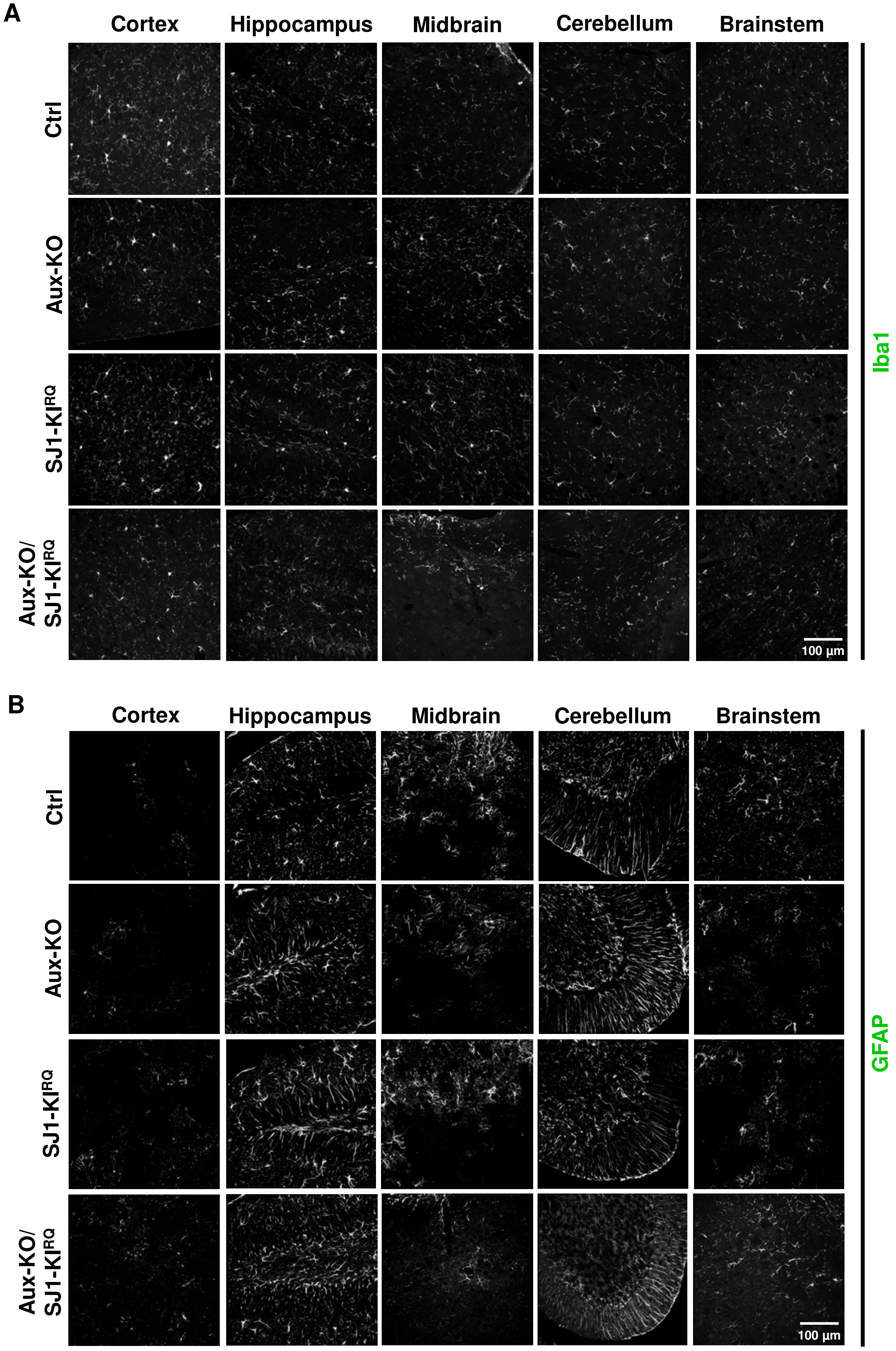


**Supplementary Fig 3**

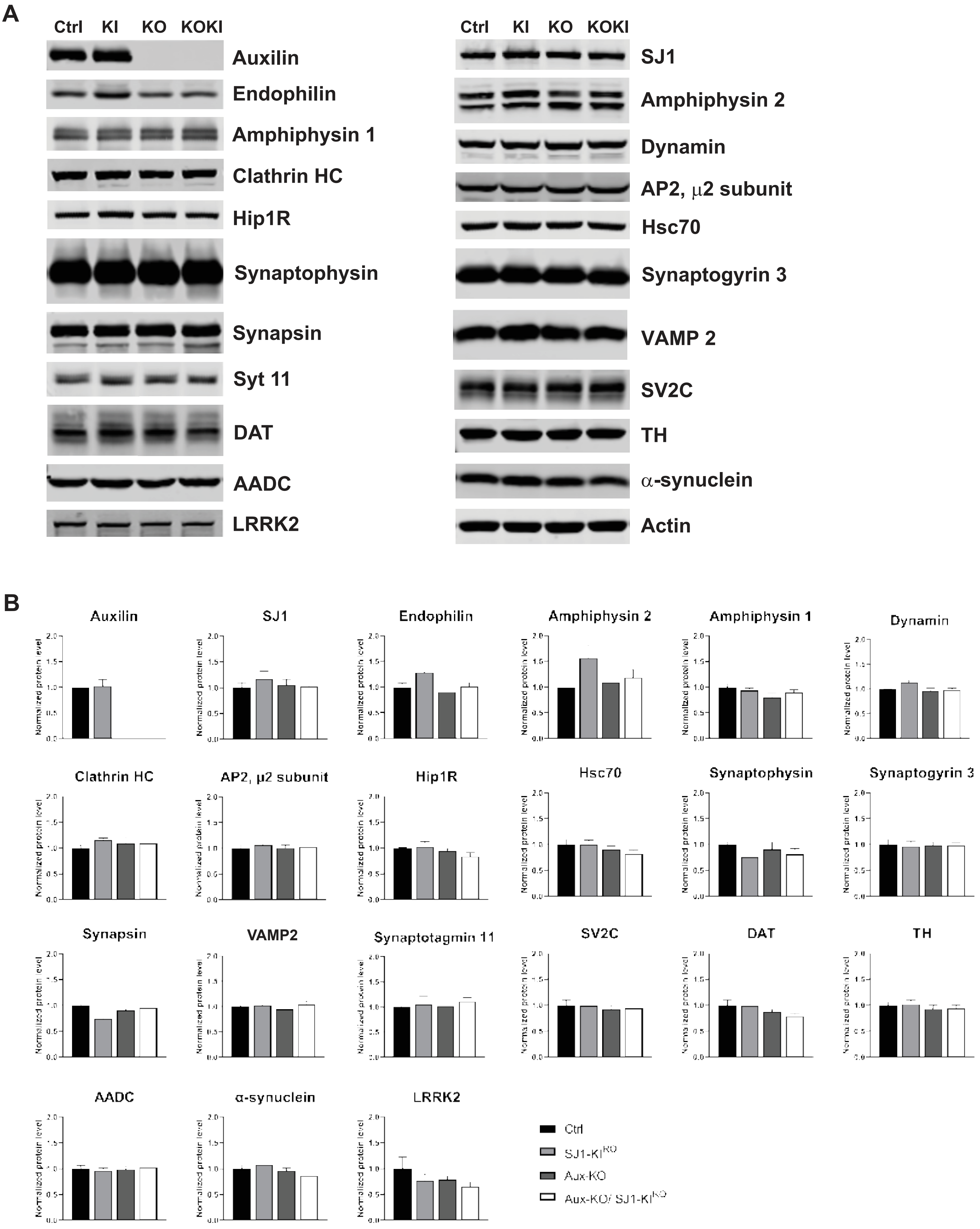


Supplementary Fig 4

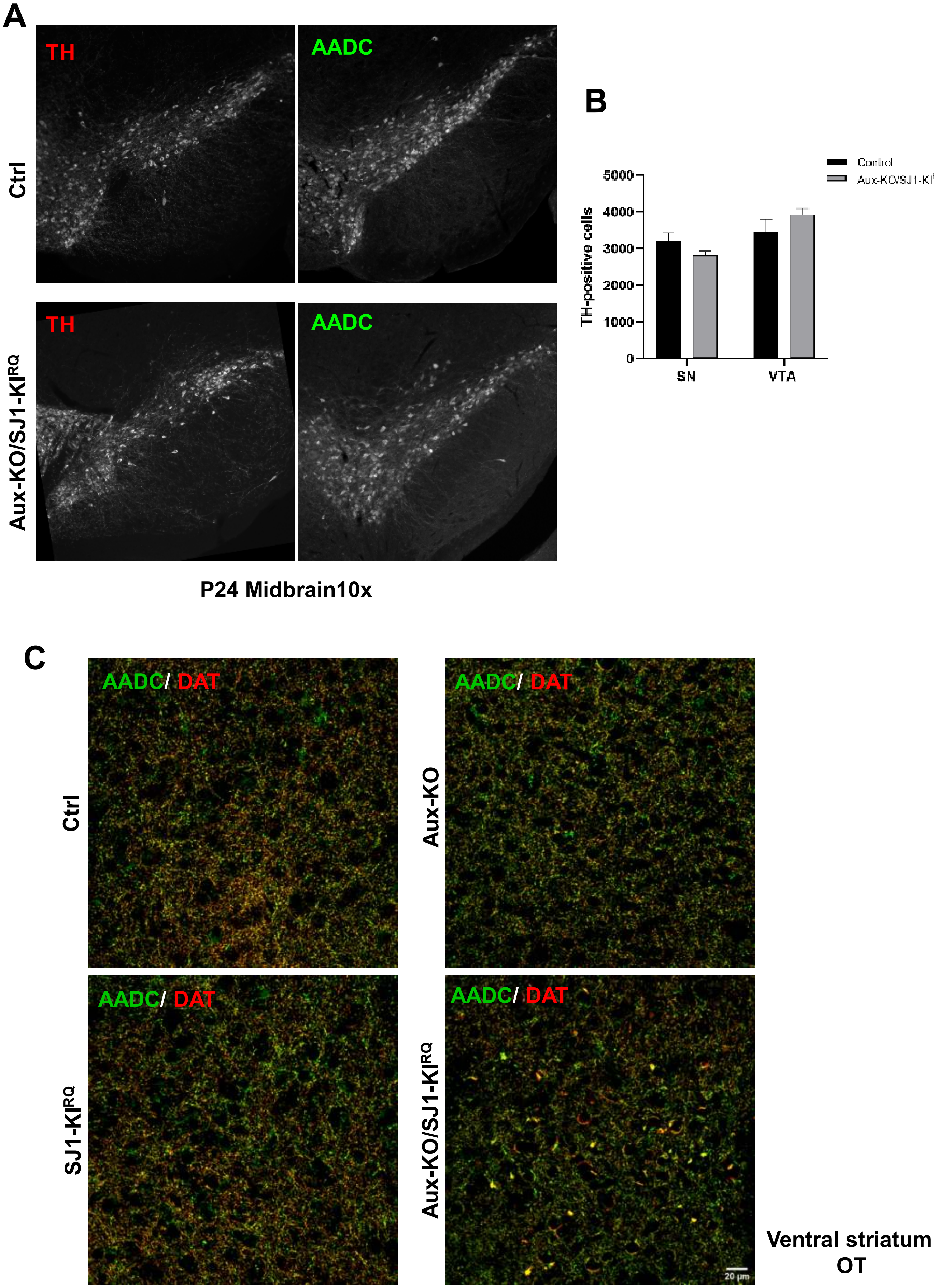


**Supplementary Fig 5**

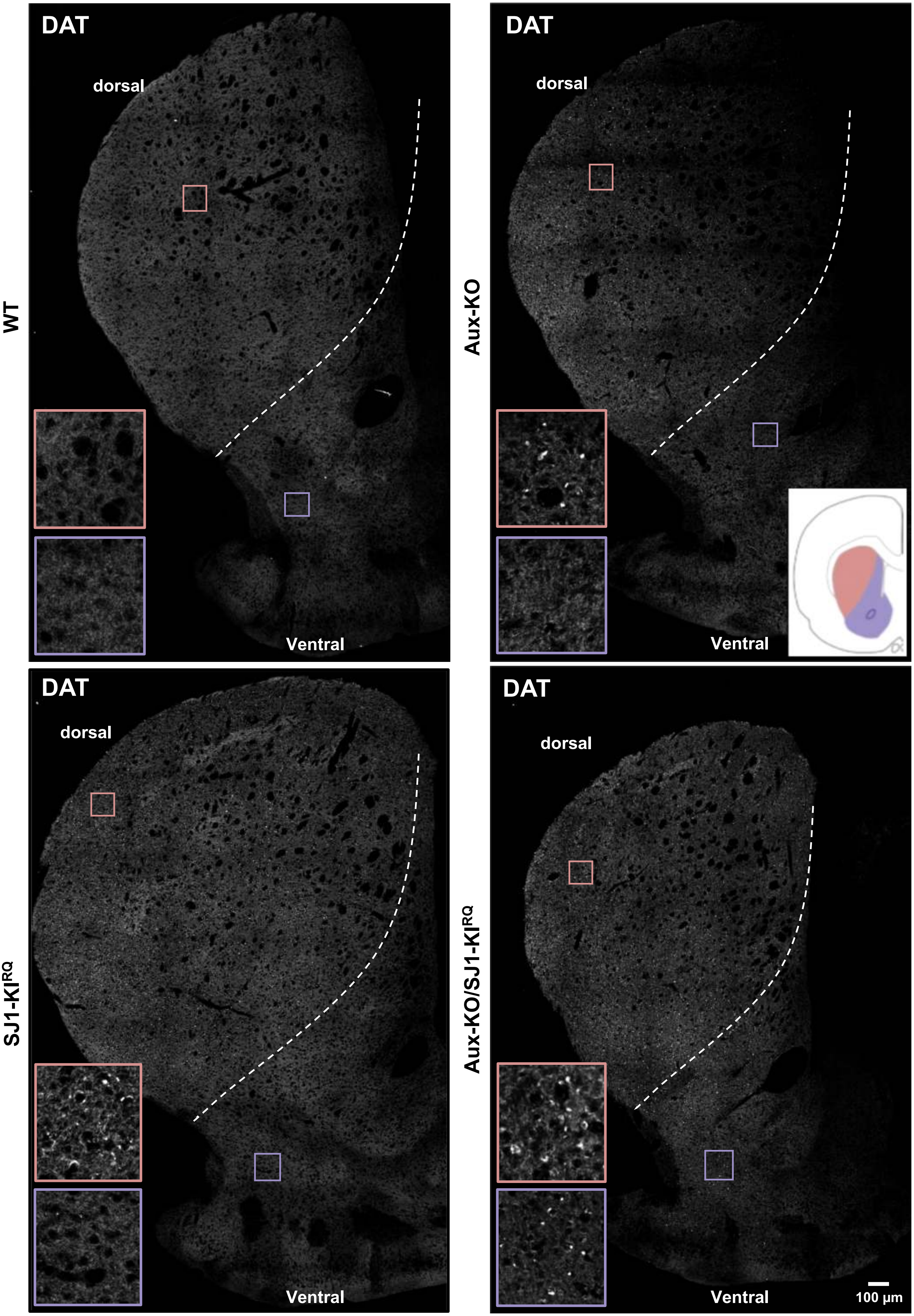


Supplementary Fig 6

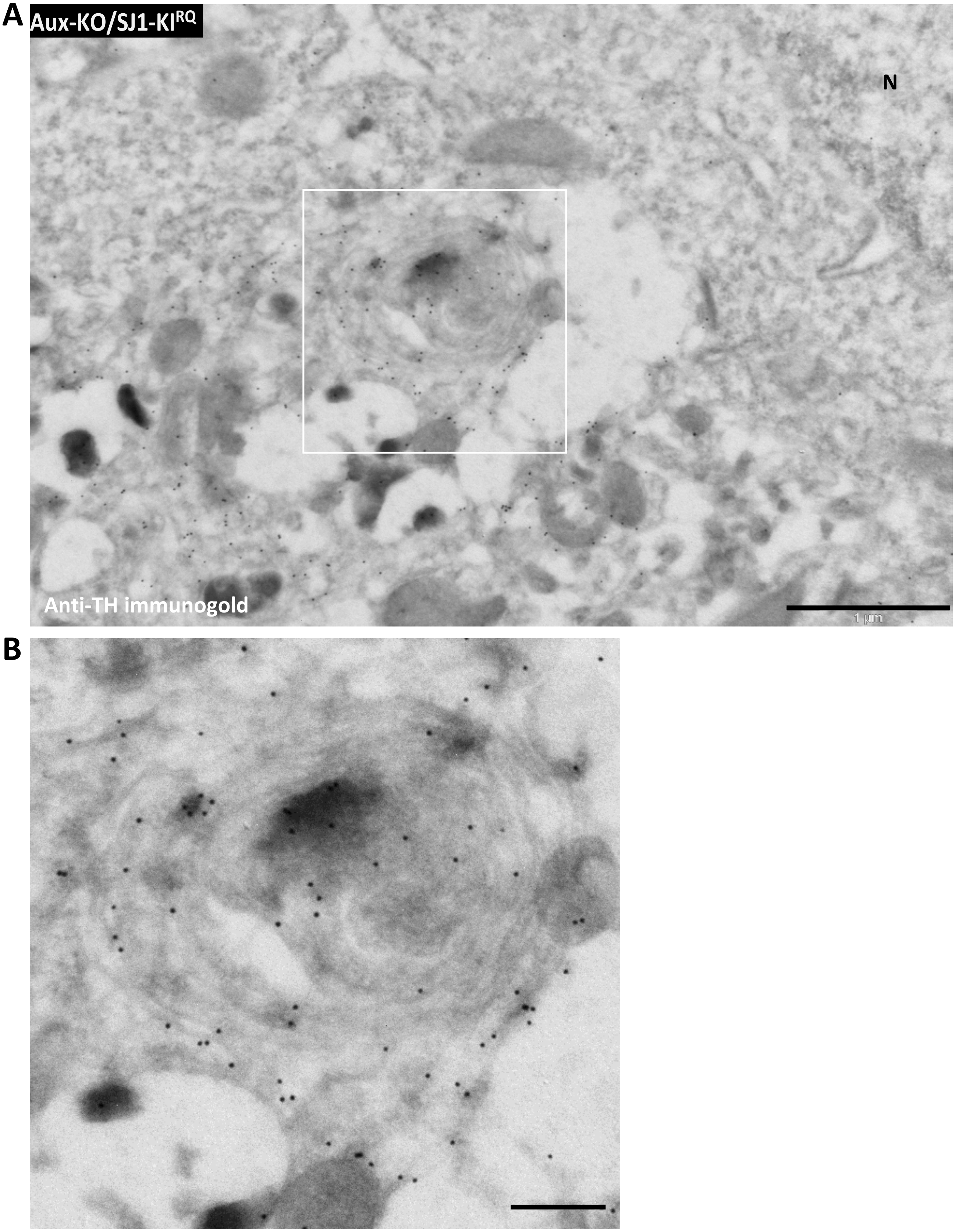


Supplementary Fig 7

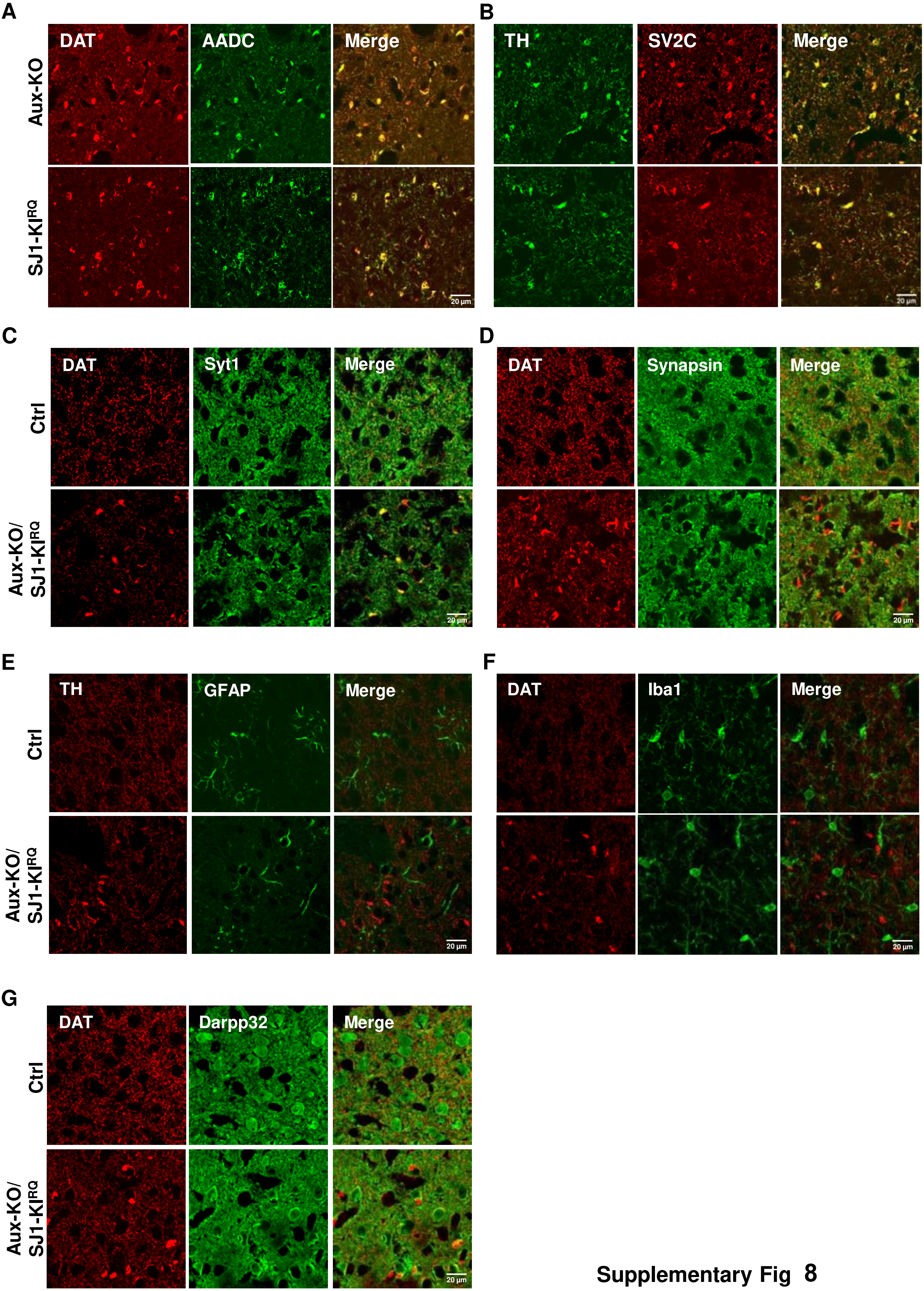


Supplementary Fig 8

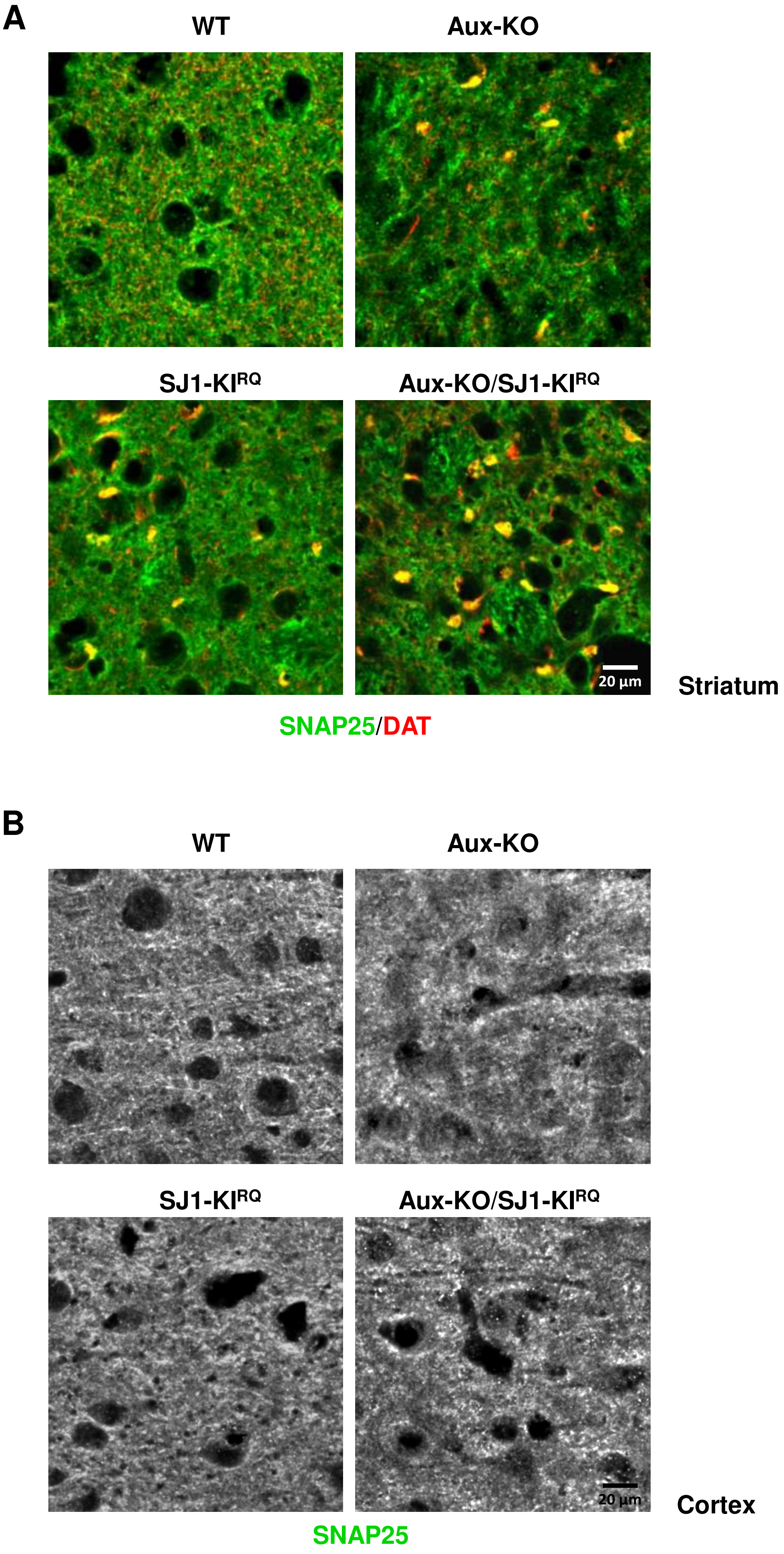


Supplementary Fig 9

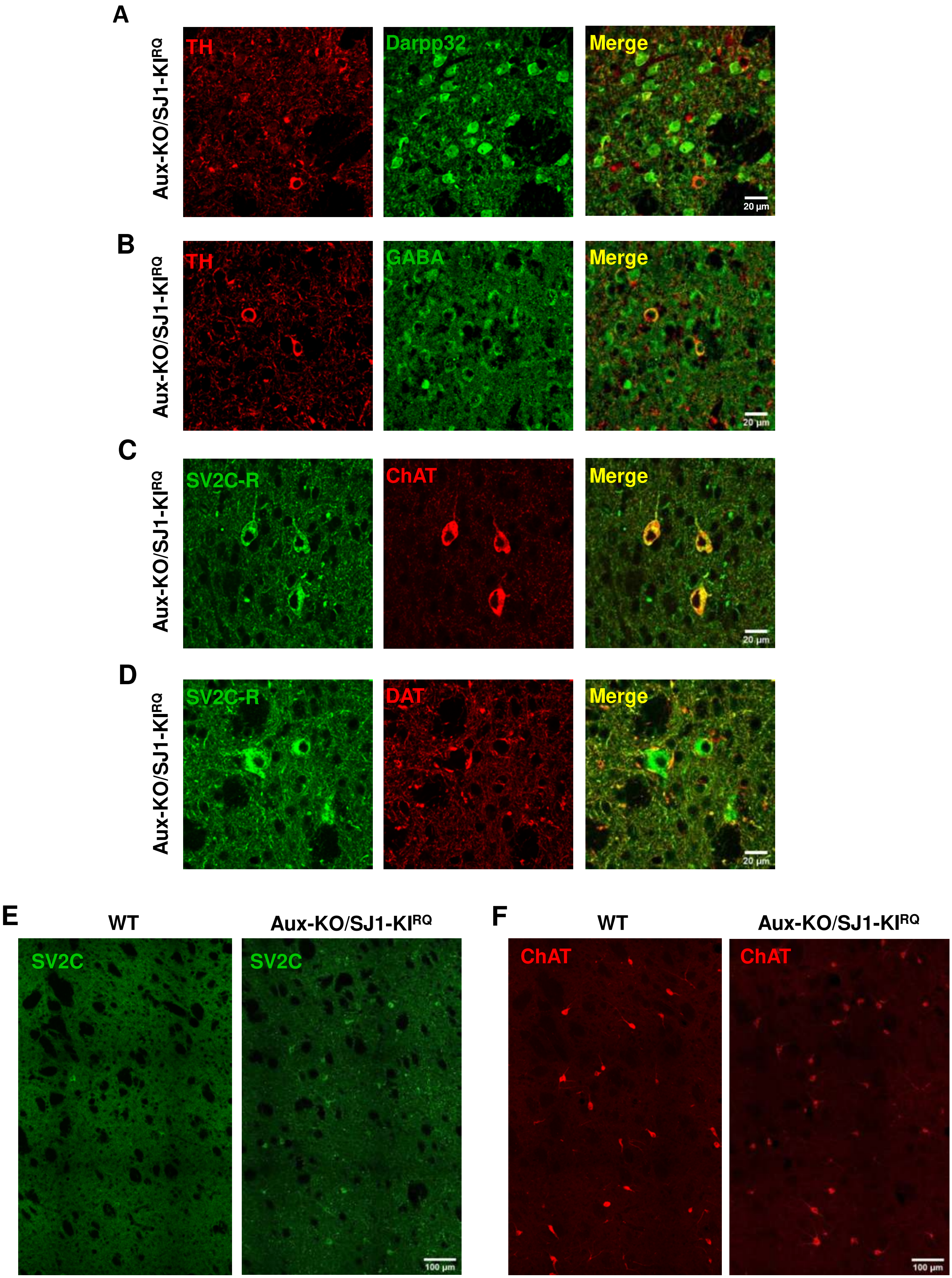


Supplementary Fig 10
